# T6SS mutants exploit itaconate to support infection of phagocytes

**DOI:** 10.64898/2026.03.11.711069

**Authors:** Ayesha Z. Beg, Blanche L. Fields, Ying-Tsun Chen, Tania Wong Fok Lung, Griffin Gowdy, Absar Talat, Asad U. Khan, Shivang S. Shah, Ian Lewis, Sebastian Riquelme, Alice Prince

**Affiliations:** Department of Pediatrics, Columbia University, New York, NY, USA; Department of Microbiology, Biochemistry and Molecular Genetics, Rutgers New Jersey Medical School, Newark, NJ, USA; Center for Immunity and Inflammation, Rutgers New Jersey Medical School, Newark, NJ, USA; Antimicrobial Resistance Lab, Interdisciplinary Biotechnology Unit, AM University, 202002, India; Bioinformatics and Computational Biology Centre of DBT Govt. of India, Interdisciplinary Biotechnology Unit, AM University, 202002, India; Department of Biological Sciences, University of Calgary, Calgary, T2N 1N4, Canada

## Abstract

*Pseudomonas aeruginosa* is a major cause of persistent pneumonias that are not readily cleared by seemingly appropriate antimicrobial therapy. We identified a reservoir of *P. aeruginosa* variants lacking expression of the H3-T6SS in patients with chronic but not acute pneumonia. A PAO1 ΔH3-T6 mutant caused increased infection in the murine lung as compared to the wild-type strain. The ΔH3 mutants exhibited increased transcription of genes involved in phagocytic uptake and respiration under conditions found in the phagolysosome, namely low O_2_, low pH and abundant itaconate. We confirmed increased intraphagocytic residence of the ΔH3 mutants and colocalization with LAMP1 within the phagolysosome of both bone marrow derived macrophages *in vitro* and in alveolar macrophages harvested directly from infected lungs. Persistence within macrophages required itaconate which preserved the viability of infected macrophages and boosted bacterial bioenergetics to optimize consumption of available carbon sources. Our findings demonstrate that selection for loss of H3-T6SS loss of function mutations promotes the metabolic versatility that enables *P. aeruginosa* to cause intractable pulmonary infection.

## Manuscript

*Pseudomonas aeruginosa* is an opportunistic pathogen that causes of fulminant sepsis as well as more indolent, but persistent pulmonary infections that are a major cause of morbidity and mortality worldwide ^1,2^. In contrast to the acute, often fatal inflammatory response activated by *P. aeruginosa* sepsis, less fulminant and more prolonged *P. aeruginosa* pulmonary infections are common even in patients with normal immune function. Clinical isolates from such infections accumulate genetic changes, especially the acquisition of antimicrobial resistance genes that reflect repeated exposure to ineffective antibiotics and the selection of variants reflecting the evolution from environmental strains into host-adapted pathogens^3^. The expression of the types 3 and 6 secretion systems (T6SS) are linked to acute versus indolent *P. aeruginosa* pulmonary infections respectively, both regulated by the GacA/RetS system^4^. Whereas the effects of the T3SS toxins in the pathogenesis of pneumonia are established, the role of the T6SS is not well defined^5^. The effectors of these secretion systems respond to specific environmental threats, directing invasive or sessile lifestyles of these pathogens as frequent causes of ventilator associated pneumonia, COPD, cystic fibrosis and infection in immunocompromised hosts ^6^.

Recruitment of phagocytes and their production of the immunometabolite itaconate is a critical component of the response to *P. aeruginosa* infection that protects the host but also shapes bacterial gene expression^7^. Bacteria are confronted by challenging conditions within the lung and especially within the phagolysosome, a compartment with low O_2_ which is rapidly converted to ROS by NADPH oxidase, low pH 4.5-5 and 10 mM concentrations of itaconate^8^.

Itaconate has antibacterial activity against some pathogens and accumulates within vacuoles in macrophages^9^. Although itaconate promotes the formation of lysosomes, an organelle critical for bacterial clearance ^10,11^ it also functions as an anti-inflammatory immunometabolite and inhibits the NLRP3 inflammasome limiting ancillary host and bacterial damage from release of IL-1β and ROS^12^. By modifying specific cysteine residues of both host and bacterial proteins, itaconation can cause gain or loss of function of diverse metabolic components^13^. In addition to its role within phagocytes, itaconate is abundant in settings of inflammation and is a major component of the airway metabolome generated by inhaled pathogens ^7^.

*P. aeruginosa* retain numerous mechanisms to sense and respond to their environment. As an oxidant, itaconate causes membrane stress for bacteria and could activate general anti- oxidant responses including the type 6 secretion system (T6SS)^14^. The T6SS is highly conserved in Gram negative pathogens in which it functions to protect bacteria from diverse environmental threats ^15,16,17^. In *P. aeruginosa* four discrete T6SSs are encoded by clusters of 15-20 genes that generate an ATP-dependent phage-like apparatus^18^. Substantial epidemiological data confirm both the conservation and heterogeneity of T6SS genes among clinical isolates of *P. aeruginosa*^19^. In contrast to the H1-T6SS which is involved primarily in interbacterial competition, the H2-and H3-T6SSs in *P. aeruginosa* have been associated with internalization into epithelial cells through the expression of the phospholipase PldB^20^ and with adaptation to low O_2_ necessary for the efficient formation of biofilms and resistance to oxidant stress ^21–24^. The participation of T6SS effectors responsive to such diverse challenges should be especially relevant to the pathogenesis of pulmonary infection, as lung microenvironments may be anaerobic, microaerobic or well oxygenated, depending upon the location in the lung itself, accumulation of phagocytes and formation of biofilm, with each setting demanding utilization of specific bacterial metabolic pathways.

We postulated that the T6SS participates in the adjustment of *P. aeruginosa* to the lung. We demonstrate that this process occurs through selection of H3-T6SS loss of function mutants which exploit the consequences of itaconate accumulation during infection by promoting metabolic adaptation to microaerobic conditions and substrates present in phagocytes and in the infected airway.

## Results

### Clinical isolates of *P. aeruginosa* acquire loss of function mutations in the H3-T6SS during chronic pulmonary infection

Genomic studies of clinical isolates have documented the conservation of the major alleles of the T6SS and highlight frequent mutations in these loci^19^. Our analysis of available genomic data bases indicated that the *P. aeruginosa* H3-T6SS loci had multiple missense and likely loss of function mutations in the essential ATPase *clp*V3 and *vgr*G3 in isolates recovered from chronic infection from cystic fibrosis patients as compared with isolates from ICU patients with acute pneumonias ^25^ (Fig. 1a-f). The apparent loss of H3-T6 function over the course of chronic infection was corroborated by several proteomic studies of *P. aeruginosa* that failed to detect the expected H3-T6SS gene products in isolates from cystic fibrosis (CF) patients in contrast to the PAO1 reference strain^26,27,28^. We found that T6SS genes are upregulated in acute pulmonary infection as compared with bacteria grown under *in vitro* conditions (Fig. 1g) and that the H3 locus, in particular, is upregulated in CF clinical isolates (Fig. 1h) but not from acute (ICU) pneumonias (Fig. 1i). The accumulation of mutations in the H3 locus over the course of chronic but not in an acute infection suggested an active selection process in response to local conditions in patients.

**Fig. 1.**
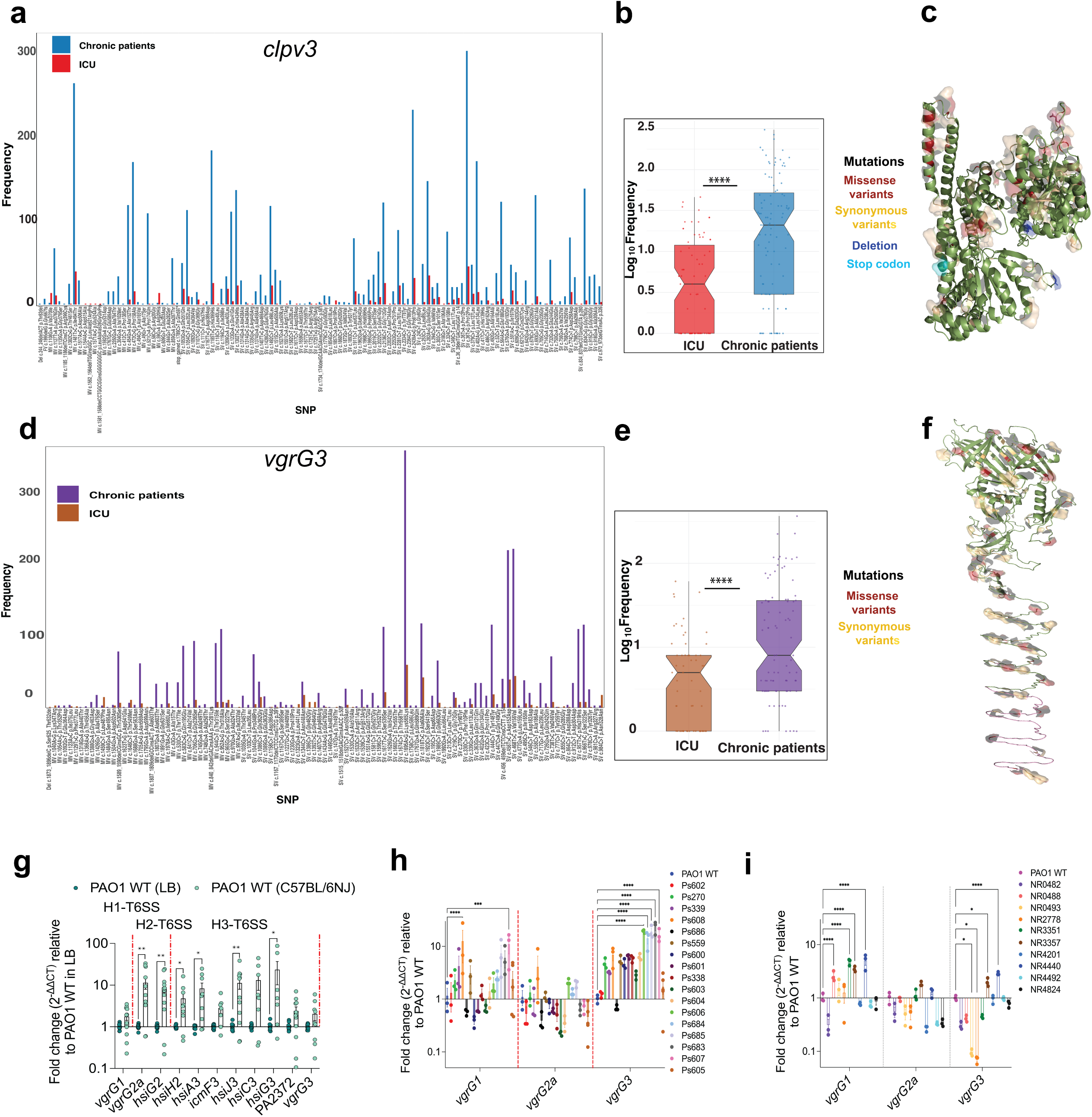
Clinical isolates accumulate loss of function mutations in the H3-T6SS locus. Relative frequency of mutations in the (a) *clpV3* and (d) *vgrG3* genes across publicly available databases of *P. aeruginosa* strains isolated from CF patients (n = 374; Bioproject accession number: PRJEB5438, PRJNA528628, PRJNA1023362, PRJNA1023843 on NCBI) and ICU patients (n = 62; ENA accession number: PRJNA629475) (SV, synonymous variant; MV, missense variant). Significant differences in total mutation frequency in (b) *clpV3* and (e) *vgrG3* between CF and acute isolates are shown by box plots. Ribbon structures of the (c) ClpV3 and (f) VgrG3 proteins highlight the sites of identified mutations. (g) In vivo expression of T6SS loci in PAO1 WT extracted from the lungs of C57BL/6NJ mice at 16 hpi relative to PAO1 WT grown in LB media, as determined by RT-qPCR (n = 3, 9 mice per group). Expression of H1 (*vgrG1*), H2 (*vgrG2a*), and H3 (*vgrG3*) loci in (h) CF and (i) ICU isolates from Columbia University relative to PAO1 WT grown in LB media, determined by RT-qPCR (n = 3). Data are presented as mean ± s.e.m. Statistical significance was assessed using two-way ANOVA with Šídák’s multiple comparisons test (g), Dunnett’s multiple-comparisons test (h, i). For (b) and (e), the Shapiro–Wilk normality test indicated that the data were not normally distributed, and significance was determined using the Wilcoxon signed-rank test (for *clpV3*: V = 59, P = 0.00058; for *vgrG3*: V = 111, P = 0.00049). All statistical tests were two-sided. Significance is defined as *P* < 0.05; \**P* < 0.01; \*\**P* < 0.001; \*\*\**P* < 0.0001. Source data are provided as a Source Data file 1.

### Loss of the H3-T6SS promotes pneumonia

We constructed mutants in the PAO1 background for each of the three major T6SS loci (Supplementary Fig. 1a) and tested outcomes in a mouse model of acute pneumonia (18 hours) using both WT C57/Bl6 and Irg1-/- mice, unable to produce itaconate. The ΔH3-T6 mutant achieved significantly <u>greater</u> levels of infection in lung tissue as compared to WT PAO1 at 18 hours in WT mice, but without significant differences in the Irg1-/- mice (Fig. 2a, b). Infection was accompanied by modestly increased production of proinflammatory cytokines commensurate with the increased bacterial load (Fig. 2c). The immune cell response to infection differed minimally between the bacterial strains (Fig. 2d, e). We confirmed that the infection phenotypes observed were due to loss of H3 function and not compensatory upregulation by the divergently regulated T3SS or H1-/H2-T6SS ^29^ (Supplementary Fig. 2a, b). In addition, equivalent bacterial loads and immune responses were found in mice infected with ΔH1, ΔH2-T6SS mutants or WT PAO1 at 18 hours of infection (Supplementary Fig. 2c-g). Thus, <u>loss</u> of H3-T6SS function specifically enhanced the ability of PAO1 to cause infection in WT mice in a model of acute pneumonia.

**Fig. 2.**
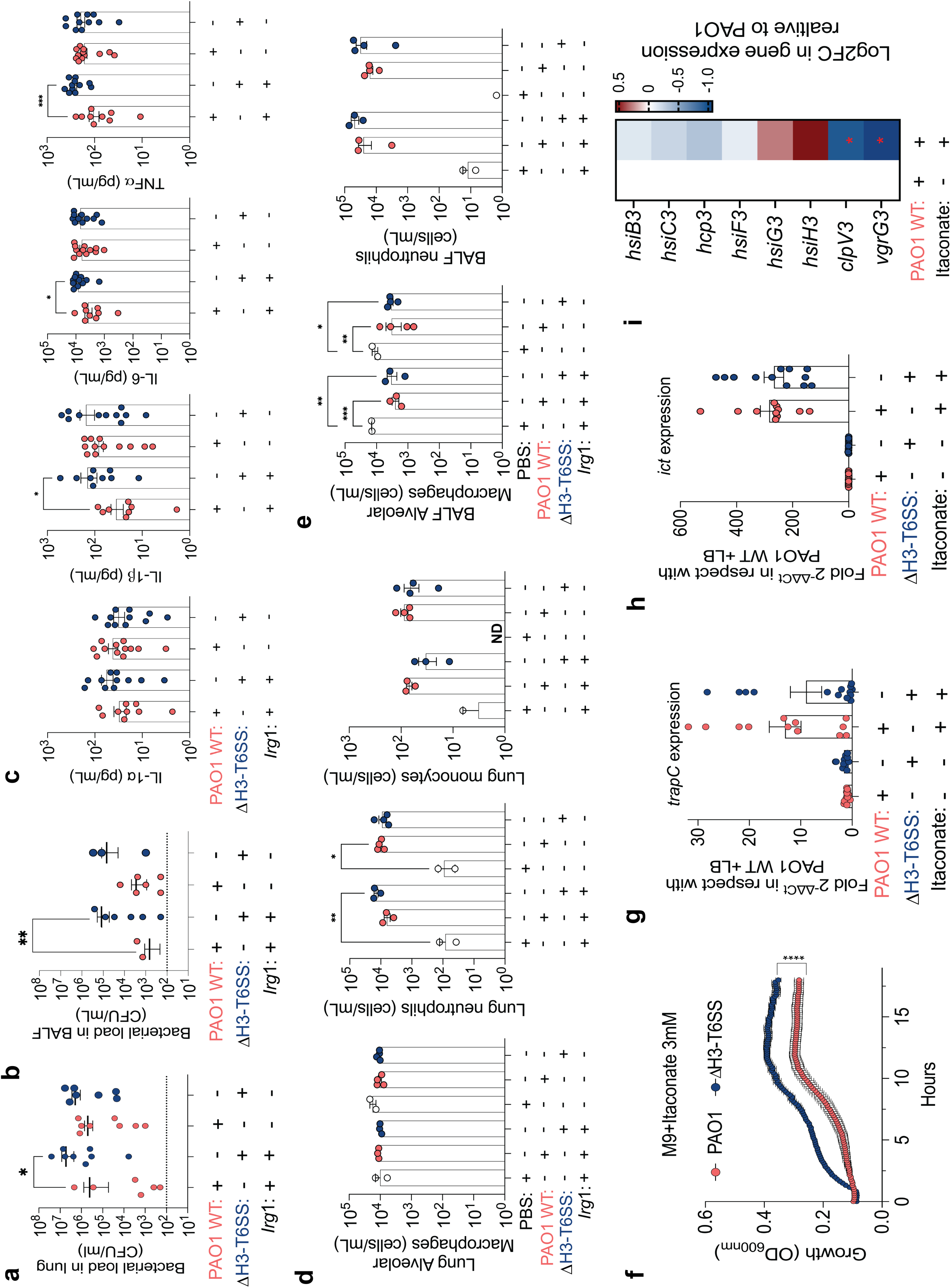
Loss of the H3-T6SS promotes pneumonia and the effects of itaconate. (a–e) Characteristics of acute pulmonary infection in C57BL/6NJ mice following intranasal inoculation with PAO1 WT or ΔH3-T6SS strains at 16 hpi. Bacterial burden in the (a) lung and (b) BALF (n = 3, 6–8 mice per group). (c) Cytokine levels (n = 3, 9–12 mice per group). Immune cell populations in the (d) lung and (e) BALF (n = 3, 3–4 mice per group). (f) Growth curves of PAO1 WT or ΔH3-T6SS strains in M9 minimal media with 3 mM itaconate as the sole carbon source (n = 3). Expression of (h) transporter (*trapC*) and (g) itaconate-metabolizing (*ict*) genes in ΔH3-T6SS relative to PAO1 WT grown in LB ± itaconate, as determined by RT-qPCR (n = 12). (i) Expression of H3-T6SS genes in PAO1 WT in response to itaconate, as determined by bulk RNA-seq (n = 2). Data are presented as mean ± s.e.m. Statistical significance was assessed using the Mann–Whitney U test (a, b, f), one-way ANOVA with Tukey’s multiple-comparisons test (c-e), and the Wald *t*-test (i). All statistical tests were two-sided. Significance is defined as *P* < 0.05; \**P* < 0.01; \*\**P* < 0.001; \*\*\**P* < 0.0001. Source data are provided as a Source Data file 2.

### Itaconate impacts the properties of the ΔH3-T6SS mutant

It appeared that host itaconate, although not critical in host defense in this acute infection model, might contribute to the increased bacterial load associated with the ΔH3-T6 mutants (Fig. 2a). *In vitro*, we found increased rates of ΔH3 growth as compared with a PAO1 WT control in physiological 3.0 mM itaconate (Fig. 2f). Appreciating that *P. aeruginosa* can metabolize itaconate, we established that both PAO1 and the ΔH3 mutant similarly increased expression of *trapC* and *ict* involved in itaconate transport and metabolism^30^ (Fig. 2g, h), indicating that the H3-T6SS is not necessary to sense itaconate as either an oxidant or substrate. We found that itaconate suppressed expression of multiple genes in the H3-T6 locus including the essential *vgrG3* and *clpv3*, an effect that could minimize differences between the WT and ΔH3-T6 strains in this model of infection (Fig. 2i). We next addressed the biological significance of itaconate on *P. aeruginosa* gene expression since it is such a prominent component of the phagocyte, phagolysosome and the infected airway.

### Itaconate promotes the uptake and viability of ΔH3-T6SS mutants in microaerobic conditions

We performed a bulk transcriptomic analysis of PAO1or ΔH3 grown in LB media with or without itaconate to determine how this host immunometabolite might impact bacterial adaptation to the lung (Fig. 3a, Supplementary Table 1). The transcriptomic data, although obtained from *in vitro* cultures grown in room air, revealed the upregulation of genes in the ΔH3 mutant in the presence of itaconate that are expected to be expressed under conditions of low O_2_ (Fig. 3a, b). One of the most highly upregulated ΔH3 genes in itaconate was *anv*M, a gene that has been associated with increased phagocytic uptake^31^. AnvM itself directly interacts with the global O_2_ sensor Anr^32^, which directs expression of Mhr (microoxic hemerythrin)^21^. Mhr interacts with cbb3-Cco oxidases which function in microaerobic conditions^33^. The O_2_ that is associated with Mhr is transported into the cell via OprG. Both Mhr and OprG are secreted by the H2-T6SS^33^ and are significantly upregulated in the ΔH3 mutant in itaconate (Fig. 3c). AnvM also interacts with MvfR (or PqsR) a part of the quinolone quorum sensing system which regulates numerous genes implicated in growth in limited O_2_ and in biofilm formation^21^. PQSR induces MvfR and itself is upregulated by PhrS, a base-pairing sRNA that operates in low O_2_ and is significantly increased in the ΔH3 mutant^34^ (Fig. 3b). Also upregulated in the mutant in itaconate and under Anr regulation are the genes encoding HemN and Dnr, which are predicted to be modified by itaconate^13,35^. We confirmed the upregulation of *anvM* and the cytochrome oxidase *cco*N2 in the ΔH3 mutant under microaerobic conditions, as well as the H2-T6SS gene *mhr*, confirming previous reports^21,33^ (Fig. 3c). The physiological significance of the transcriptomic responses was confirmed by documenting increased growth of the ΔH3 mutant as compared with PAO1 under microaerobic conditions (1% O_2_) and anaerobic conditions in the presence of itaconate and *at* low pH (Fig. 3d, e) (Supplementary Fig. 3a). We did not observe any effects of the ΔH3 mutation on biofilm formation (Supplementary Fig. 3b).

**Fig. 3.**
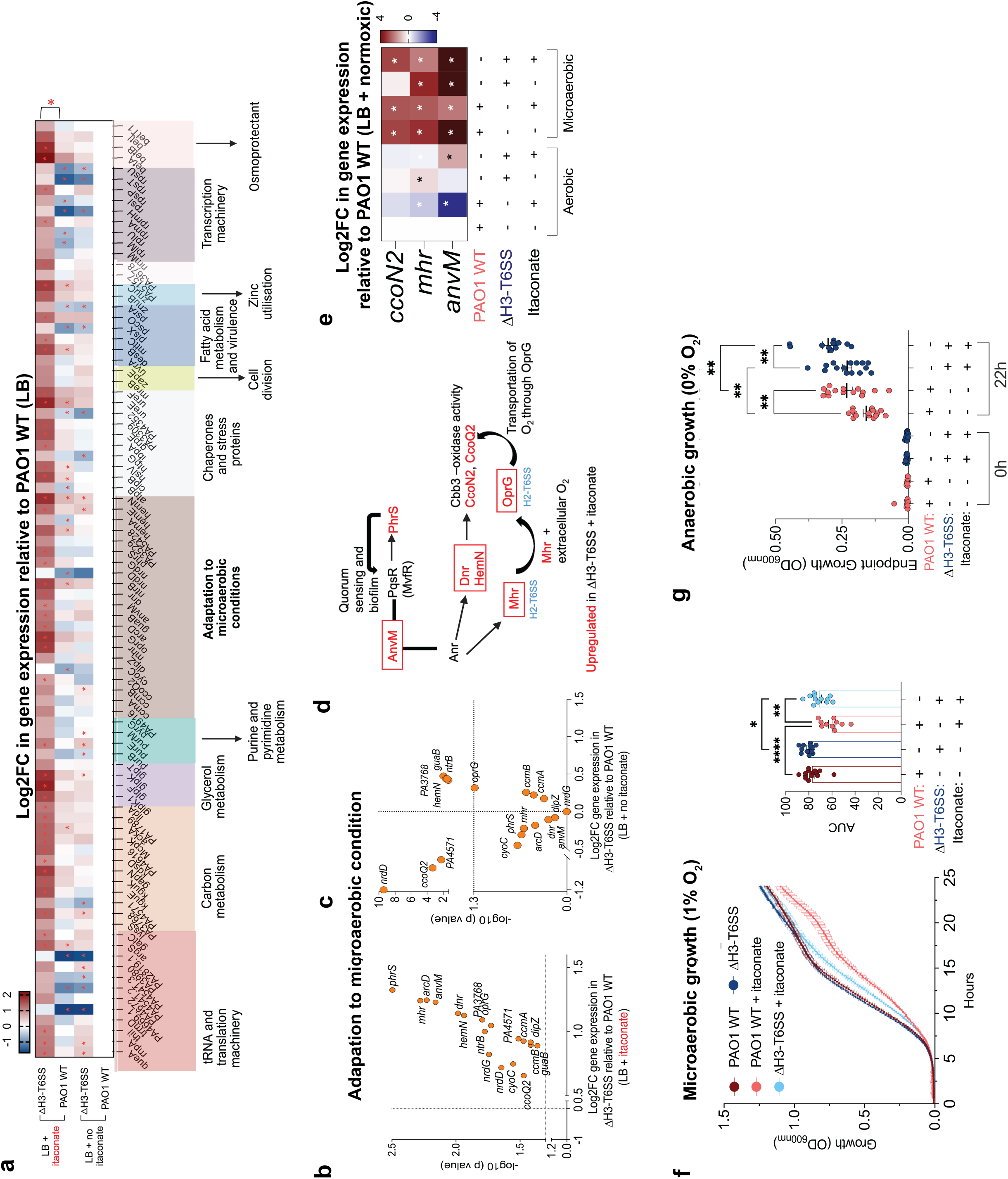
Itaconate promotes adaptation of H3-T6SS mutants to a microaerobic environment. (a) Differential expression of genes in ΔH3-T6SS or PAO1 WT grown in LB +/- itaconate normalized to PAO1 WT in LB alone, shown as a heatmap from bulk RNA-seq analysis (n = 2) (* significant). Expression of genes associated with microaerobic adaptation in ΔH3-T6SS relative to PAO1 WT in the (b) presence or (c) absence of itaconate as determined by bulk RNA-seq. (d) Schematic representation of itaconate dependent upregulation of genes associated with microaerobic pathways in ΔH3-T6SS. (e) Expression of genes associated with microaerobic adaptation (*ccoN2*, *mhr, anvM*) in normoxic or microaerobic (1% O_2_) conditions in ΔH3-T6SS or PAO1 WT grown in LB +/- itaconate as determined by RT-qPCR quantification(n = 6). (f) Growth curves of PAO1 WT or ΔH3-T6SS strains in LB +/- itaconate under 1% oxygen conditions (n = 3). (g) Growth (OD_600nm_) of PAO1 WT or ΔH3-T6SS strains cultured in LB +/- itaconate under anaerobic (0% O_2_) conditions at 22h. Data are presented as mean ± s.e.m. Statistical significance was assessed by one-way ANOVA with uncorrected Fisher’s LSD (e), Tukey’s multiple comparisons test (f, g), and Wald-*t* test (a, b, c). Schematics in (d) is created using BioRender. Significance is defined as *P < 0.05; **P < 0.01; ***P < 0.001; ****P < 0.0001. All statistical tests are two-sided. Source data are provided as a Source Data file 3.

Thus, loss of H3 function mutants are adapted to an environment replete with itaconate but with limited O_2_ availability and low pH, the conditions in the phagolysosome.

### Increased accumulation of ΔH3-T6SS mutants in macrophages is dependent upon itaconate

To directly test the central hypothesis that ΔH3-T6 mutants are adapted to conditions within the phagocyte, we quantified bacterial survival within bone marrow derived macrophages from mice (BMDMs). We found significantly greater numbers of viable ΔH3 mutants than PAO1 at 1.5 hours following infection of WT BMDMs (Fig. 4a) and no differences in the viability of the macrophages infected with either bacterial strain (Fig. 4b). In contrast, there was minimal retention of either bacterial strain in BMDMs harvested from Irg1-/- mice (Fig. 4a) and viability of infected Irg1-/- BMDMs was less than 50% at 1.5 hours of infection (Fig. 4b), suggesting a requirement for itaconate to maintain phagocyte viability. Itaconate was not necessary to fuel *P. aeruginosa* survival in macrophages as Δ*ict* mutants (either in PAO1 or a Pa 686 clinical strain) unable to consume and thus deplete itaconate sustained increased levels of intracellular infection (Fig 4. c, d). Rates of clearance of ΔH3 mutants and PAO1 were similar *in vitro* (Fig. 4f). Thus, itaconate promotes intracellular residence of the bacteria, but not by functioning as a carbon source.

**Fig. 4.**
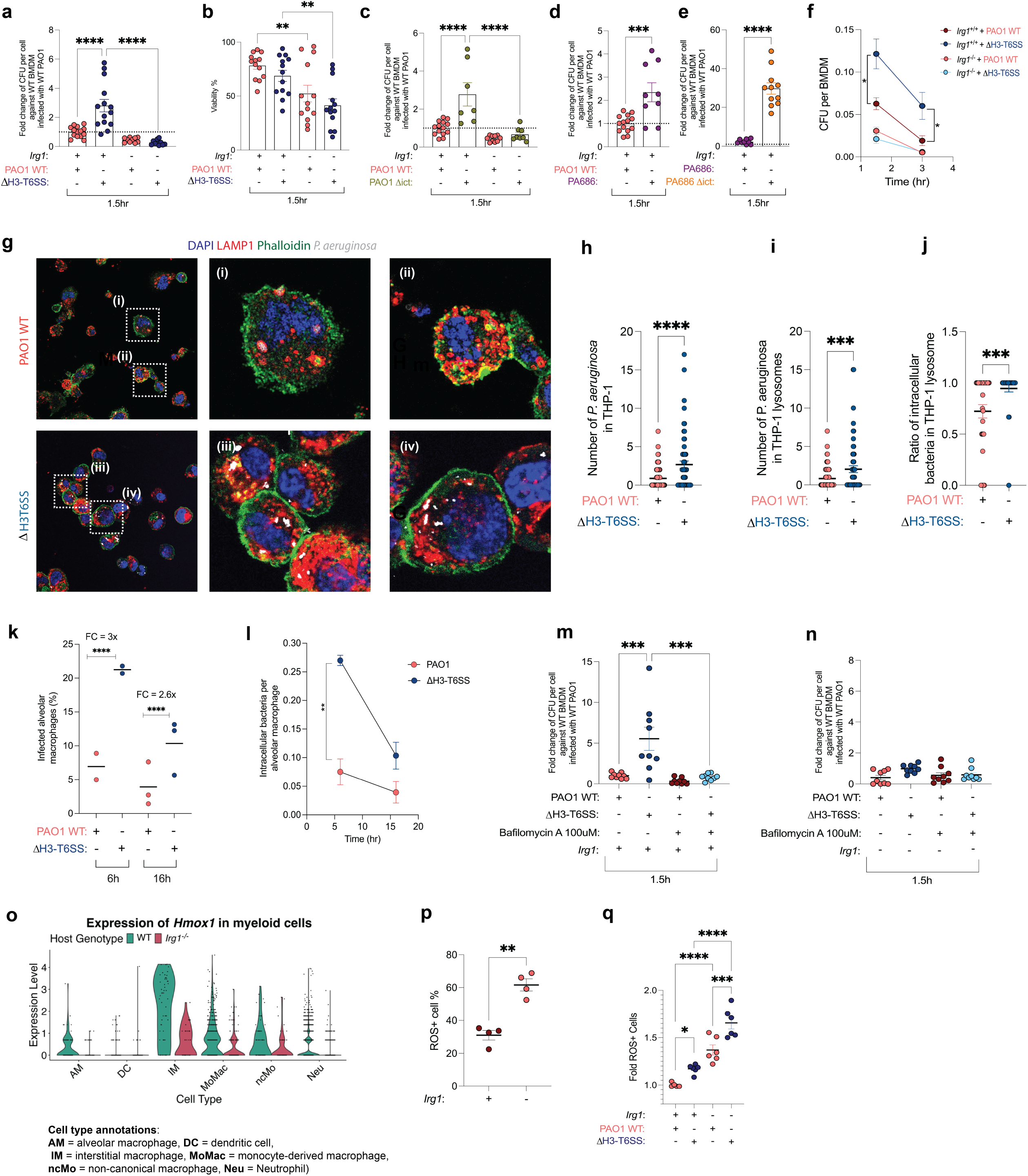
Relative levels of macrophage infection as a function of *Irg*1 expression. (a-c, e) Intracellular bacterial burden in C57BL/6NJ WT or *Irg1*⁻/⁻ BMDMs was quantified by gentamicin protection assay: (a) ΔH3-T6SS vs PAO1 WT, (c) PAO1 WT vs Δ*ict*, (d) PAO1 WT vs CF isolate 686, and (e) PAO1 WT vs CF isolate 686 Δ*ict* (n = 6-9). (b) BMDM survival following infection. (f) Kinetics of intraphagocytic uptake and survival of ΔH3-T6SS or PAO1 WT in BMDMs at 1.5 and 3 h post infection. (g) Intraphagocytic colocalization of(i-ii) PAO1 or (iii-iv) ΔH3 with lysosome marker, LAMP1. (h) Total intracellular and (i) lysosome-localized intracellular bacteria in THP-1 cells. (j) Ratio of lysosome-localized to total intracellular ΔH3-T6SS or PAO1 WT bacteria in THP-1 cells. (k) Percentage of infected alveolar macrophages (AMs). Bars indicate the mean and dots represent biological replicates; significance was assessed by Fisher’s exact test on pooled cell counts. FC denotes the fold-change in infection frequency between ΔH3-T6SS and PAO1 WT at each time point. (l) Kinetics of intracellular bacteria per AM at 6 and 16 hpi. Intracellular bacterial burden quantified by gentamicin protection assay ± bafilomycin in (m) WT and (n) *Irg1*⁻/⁻ BMDMs infected with ΔH3-T6SS or PAO1 WT (n =9). (o) scRNA-seq based expression of *Hmox*1 in myeloid cell populations in C57BL/6NJ WT or *Irg1*⁻/⁻ mice infected with *P. aeruginosa*. (p) Baseline (uninfected) percentage of ROS-positive and (q) infection-induced fold change in ROS-positive WT or *Irg1*⁻/⁻ BMDMs (n = 3-6). Data are presented as mean ± s.e.m. Statistical significance was assessed by one-way ANOVA with Tukey’s multiple comparisons test (f, i, j, m, n), Uncorrected Fisher’s LSD (a-e, q), two-way ANOVA with Šídák post-hoc test (l; interaction P = 0.0227, time P = 0.0033, strain P = 0.0009), unpaired t-test with Welch’s correction (p), and Mann-Whitney U test (h). Significance is defined as *P < 0.05; **P < 0.01; ***P < 0.001; ****P < 0.0001. All statistical tests are two-sided. Source data are provided as a Source Data file 4.

Based on our transcriptomic data the increased intraphagocytic residence of the ΔH3- T6 mutants seemed likely to be due to AnvM associated uptake and enhanced intracellular survival of the mutants, due to bacterial adaptation to the conditions within the phagolysosome. Using the human THP-1 macrophage cell line we found increased internalization of the ΔH3 mutant and a significantly greater fraction of the intracellular bacteria co-localizing with a LAMP-1 marker for the phagolysosome (Fig. 4g-j) (Supplementary Fig. 4a). To confirm the phagolysosomal location of the ΔH3-mutants *in vivo*, we isolated alveolar macrophages from BAL fluid from infected mice at 6 and 16 hours post infection, fixed and stained them for both LAMP-1 and *P. aeruginosa* (Fig. 4k) (Supplementary Fig. 4b-d). At 6 hours post infection ΔH3-T6 bacteria were 3-fold more abundant than WT PAO1 within alveolar

macrophages colocalizing with LAMP-1. At 16 h post infection, ΔH3-T6 mutants remained 2.6- fold more abundant than WT PAO1 despite increased bacterial clearance (Fig. 4 k, l).

We attributed the greater intraphagocytic infection by the ΔH3 mutant to increased uptake and survival within the phagolysosome as compared with PAO1. Although increased phagocytosis of bacteria is predicted to increase bacterial clearance, the metabolic adaptation of the ΔH3 mutants to conditions within the phagolysosome would explain increased rates of infection both *in vivo* (Fig. 2a, Fig. 4k) and *in vitro* (Fig. 4a). To confirm the increased residence of the ΔH3 mutant within the phagolysosome we treated WT and Irg1-/- BMDMs with the vATPase inhibitor bafilomycin A that prevents acidification of the phagolysosome^36^. Although bafilomycin treated macrophages should retain greater numbers of bacteria, we observed that by neutralizing the phagolysosome, we removed the conditions selectively favorable for the ΔH3 mutant and eliminated its advantage in intraphagolysosomal retention that was observed in control phagocytes (Fig. 4m, n). These phagocytosis studies suggest that the ΔH3 mutant is preferentially retained in the phagolysosome where it is adapted to low pH, low O_2_ and high itaconate, even if only transiently.

We noted in the phagocytosis assays that macrophages lacking itaconate, both bafilomycin treated as well as untreated controls, retained few bacteria, indicating an unexpected role for itaconate in permitting macrophage infection. Bacterial infection stimulates macrophage production of itaconate which triggers anti-oxidant responses through the Keap1- Nrf2 pathway^37^. Itaconate by itself did not activate expression of anti-oxidant genes in *P. aeruginosa* (Supplementary Fig. 4e). Host protection from oxidant stress is mediated by the Nrf2 dependent enzyme Hmox1^38^. We found that *Irg1* is critical in regulating *Hmox-1* expression in myeloid cells during PAO1 infection (Fig. 4o), which helps to explain why *Irg1* is important in sustaining macrophage survival during infection. We established that in the basal state there was significantly increased production of ROS in *Irg1*-/- BMDMs (Fig. 4p) and upon infection, a greater proportion of the *Irg1-/-* cells produced ROS than the WT controls (Fig. 4q). Of note the ΔH3 mutant generated more ROS than PAO1(Fig. 4q) as well as greater amounts of IL-1α, IL-1β and IL-18 (Supplementary Fig. 4f-h), indicating that increased intracellular retention is not due to decreased immunogenicity of the ΔH3 mutant. Thus, itaconate as a host protectant is especially important in enabling the intraphagocytic retention of the more immune stimulatory ΔH3 mutant.

### Itaconate amplifies the metabolic activity of the ΔH3 mutant

Persistence of the ΔH3 mutants within phagocytes implies bioenergetic capabilities within this challenging milieu and not just transitory passage of organisms continuing to use aerobic respiration. Itaconate can have either gain or loss of function consequences on specific host and bacterial metabolic targets. We performed a direct comparison of intracellular metabolites generated by bacteria grown in the presence or absence of itaconate *in vitro* (Fig. 5a, b, Supplementary Table 2). We observed that in the presence of itaconate the ΔH3 mutant had increased accumulation of lactate, as well as NAD and FAD, key molecules essential for shuttling electrons to generate ATP. The specific pathways contributing to bacterial metabolism in the presence of itaconate were suggested by the bulk transcriptional data (Fig. 3a). We had noted upregulation of *lldA* which enables *P. aeruginosa* to metabolize lactate to pyruvate generating NADH (Fig. 5c). Lactate is a substrate for *P. aeruginosa* within phagocytes ^39^ and a major component of the phagocyte metabolic response to infection, previously observed in transcriptomic studies of CF infections ^40,41^.

**Fig. 5.**
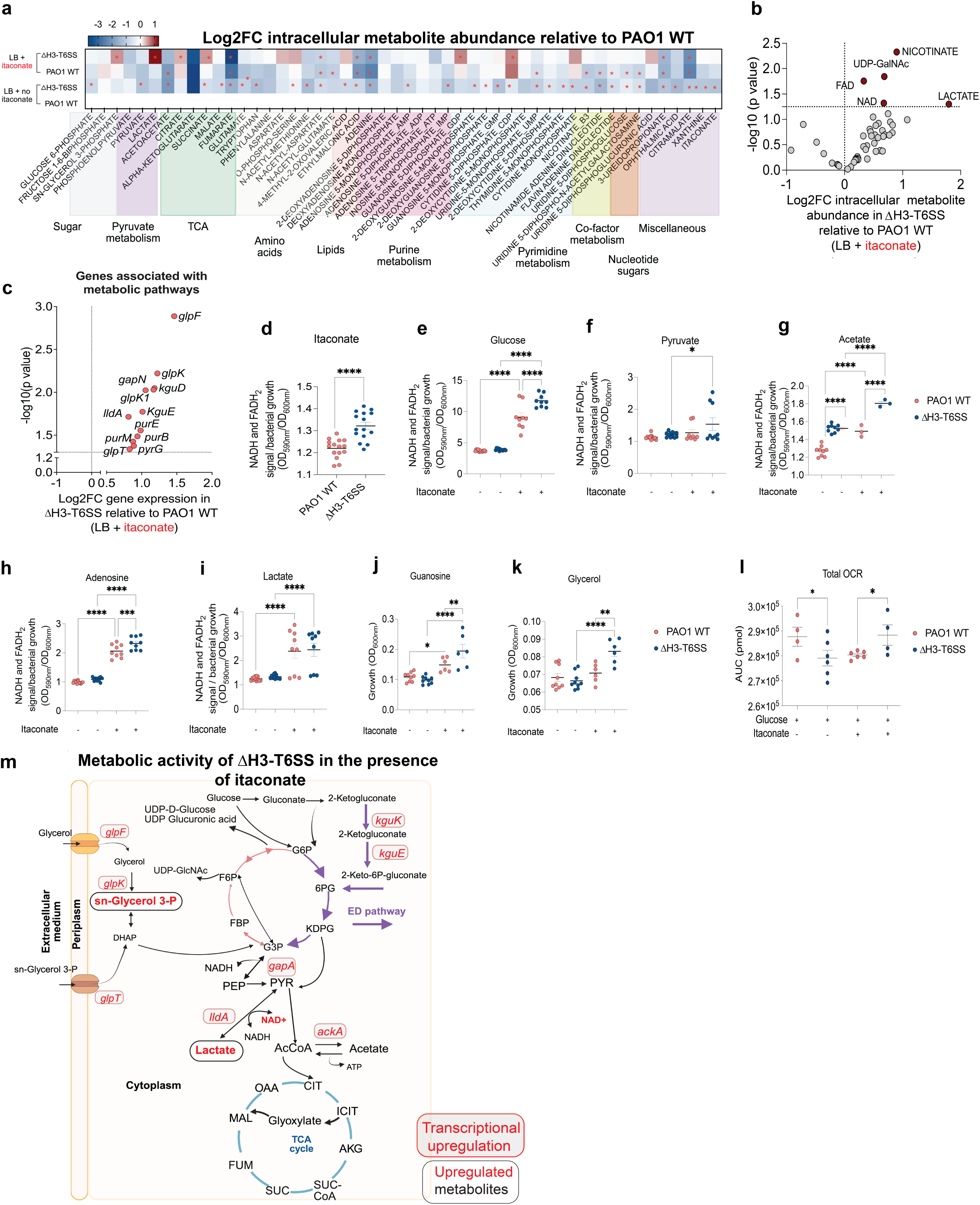
Itaconate alters metabolic activity of ΔH3-T6SS mutants of *P. aeruginosa*. (a) Accumulation of intracellular metabolites in PAO1 WT or ΔH3-T6SS in LB +/- itaconate normalized to PAO1 WT in LB alone, displayed as a heat map (n = 3) (* significant). (b) Intracellular metabolite abundance in ΔH3-T6SS relative to PAO1 WT grown in LB+ itaconate as shown in a volcano plot. Grey dots represent all detected metabolites; maroon dots represent significantly abundant metabolites. Y-axis: logarithmic scale, cut off set on Y-axis: -log10 (*p* value) ≥ 1.3; cut off set on X-axis Log2FC (Fold change) > 1). (c) Differentially expressed metabolic genes in ΔH3-T6SS relative to PAO1 WT in the presence of itaconate as shown in the volcano plot. (d-i), Generation of NADH or FADH_2_ colorimetric signal normalized to growth (n=3) for (d) itaconate, (e) glucose, (f) pyruvate, (g) acetate, (h) adenosine, and (i) lactate. Growth (OD_600_) of ΔH3-T6SS or PAO1 WT in (j) guanosine or (k) glycerol +/- itaconate. (l) Total oxygen consumption rate (OCR) (pmol) of WT vs ΔH3-T6SS PAO1 strains in glucose +/- itaconate (n = 4-6). (m) Schematic model illustrating metabolic changes of ΔH3-T6SS in response to itaconate relative to PAO1 WT, highlighting upregulated metabolic genes (red box) and significantly altered metabolites (red). Data are presented as mean ± s.e.m. Statistical significance was assessed, one-way Anova with uncorrected Fisher’s LSD (d-k, l), Mann Whitney U test (a, b), and Wald *t* test (c). Schematics in (m) is created using BioRender. Significance is defined as *P < 0.05; **P < 0.01; ***P < 0.001; ****P < 0.0001. All statistical tests are two-sided. Source data are provided as a Source Data file 5.

The transcriptomic data also suggested that the ΔH3 mutants grown in itaconate preferentially shunt glucose into glucuronates as indicated by increased expression of *kguK* and *kguE* which generate ketogluconate (Fig. 5c). This reflects use of the Entner-Doudoroff pathway^42^ a favored pathway that is promoted by itaconation of RpoN^43^. We found significant upregulation of the glycerol transporters at the level of transcription (*glyF, glyT, glypK*) consistent with the increased intracellular glycerol 3-phosphate identified in the metabolomics analysis (Fig. 5a). In addition, increased metabolism of purines by the ΔH3 mutant in itaconate was suggested by the upregulated activity of *purE* and *purM* (Fig. 5c) and consistent with reported effects of itaconate on this pathway^13^. Purines as degradation products of DNA are available within phagocytes as well as in the infected airway.

The significance of these specific substrates is their metabolism to produce FADH_2_ and NADH to generate ATP for ongoing metabolism and growth (Fig. 5b). We found that the ΔH3 mutant had significantly increased production of FADH_2_/NADH from the assimilation of itaconate alone; or plus lactate, glucose, adenosine, pyruvate or acetate (Fig. 5d-h). Itaconate plus glycerol or guanosine increased ΔH3 growth as compared with PAO1 (Fig. 5i, j). Thus, the ΔH3 mutant is primed to exploit carbon sources readily available either in phagocytes or in the infected lung, namely lactate, purines, glycerol. The overall impact of itaconate on ΔH3 metabolism was further reflected by oxygen consumption rates (OCR), a downstream readout of electron transport chain activity, which were decreased under control conditions but increased in the presence of itaconate as compared to PAO1 (Fig. 5l) (Supplementary Fig. 5a, b).

We also performed a metabolomic analysis of the broncho-alveolar lavage fluid from infected mice which reflects both host and bacterial metabolic activity (Supplementary Fig. 5c, d). We found significantly less hypoxanthine, adenosine, succinate and fumarate in the ΔH3 BAL fluid of WT mice. These findings could reflect increased purine metabolism and TCA cycle activity in the mutant. There were fewer significant differences in the airway metabolites in infections in Irg1-/- mice, highlighting the importance of host immune cell itaconate on metabolic activity during infection.

The intracellular metabolomic data, transcriptomic analysis and physiological studies each demonstrate that loss of H3-T6 function by itself has an impact on bacterial metabolism that was very substantially increased in the presence of itaconate as omnipresent *in vivo*.

These findings indicate that the prevalence of clinical isolates with loss of function mutations in the H3-T6SS reflects a substantial benefit to the pathogen in the setting of pulmonary infection; optimized metabolism of local substrates and enhanced bioenergetics that enable sequestration within phagocytes and in the airway.

## Discussion

We provide strong evidence that the T6SS is involved in the pathogenesis of *P. aeruginosa* pulmonary infection with selection for loss of function mutants over the course of chronic infection in a setting of abundant itaconate. These mutants exhibit increased expression of *anvM* which promotes phagocytic uptake along with downstream effectors that enhance bioenergetics under conditions of limited O_2_, characteristic of the phagolysosome, within biofilms and in the infected airway^21^. The organisms readily consume substrates generated by the degradation products present within the phagocytes^8^ or from the surfactant metabolite glycerol^44^. While *P. aeruginosa* is not a fully competent intracellular pathogen, as is *Coxiella* or *Mycobacteria*, like other ESKAPE pathogens ^45^, particularly *Staphylococcus aureus* ^46^, these nominally extracellular aerobic bacteria can maintain viability within the phagolysosome. This may provide a site for infection, not typically targeted by clinicians. The most common anti-*Pseudomonas* antibiotics fail to accumulate within phagocytes. Our data demonstrate the metabolic changes accompanying loss of the H3-T6SS that enable *P. aeruginosa* to persist in this challenging environment, indicating that we have not just caught them as they transit through the phagolysosome.

Host itaconate is a critical factor necessary to make the macrophage conducive for even transient bacterial residence. Fundamentally, itaconate stimulates the generation of lysosomes via TFEB expression^10^ and hence the formation of phagolysosomes critical for bacterial clearance^47^. However, the role of itaconate in promoting expression of anti-oxidants such as *Hmox1* and suppressing pyroptosis^12^ preserves macrophage viability and provides a nidus for bacterial residence. Other pathogens which stimulate itaconate production such as *Mycobacteria* or *Salmonella*^8^ also exhibit metabolic changes associated with itaconation of key enzymes^13^ and actively assume an intraphagosomal lifestyle. Our studies did not define the nature on the phagosomal structure occupied by *P. aeruginosa*, only that the majority of the ΔH3 mutants are associated with LAMP-1 positive compartments. However, the effects of baflinomycin in negating the increased intraphagocytic residence of the ΔH3 mutants, along with the transcriptomic, metabolomic and physiological data all suggest that they are adapted to the low pH, low O_2_ and high itaconate that are characteristic of the phagolysosome.

The selective expression of components of the T6SS of *P.aeruginosa* promote the overall survival of the bacterial community by regulating phenotypic heterogeneity that includes intracellular residence, biofilm formation as well as planktonic growth^48^. The H3-T6SS mutants are selected in vivo for their optimized metabolic activity in response to the phagolysosome milieu. This is quite distinct from observations with the obligate intracellular pathogen *Francisella* which uses the T6SS to break out of the phagolysosome and gain access to essential substrates in the cytosol^42^. Since *P. aeruginosa* adaptation to low O_2_ is relevant to biofilm formation^41,49^ as well as anaerobic environments^50^ the metabolic adaptive changes of the ΔH3 mutants would also enhance survival in extracellular, hypoxic areas of the lung. Our findings suggest that expression of the pathways optimized for microaerobic conditions, whether within the infected airway or the phagolysosome, are suppressed by the H3-T6SS, under oxygen-replete conditions, in the absence of itaconate, as typical for laboratory conditions, but not the infected lung. Exactly which other bacterial functions are derepressed in the absence of the H3-T6SS locus remain to be defined. Since the *in vivo* phenotype of the ΔH3-T6 mutant in the absence of itaconate is minimal, host itaconate clearly drives the selection of H3-T6SS variants *in vivo*; as it is necessary for the potentiation of the metabolic activity unleashed by the loss of H3 function as well as the preservation of macrophage viability.

The overarching goal of this project has been to determine how the T6SS contributes to the pathogenesis of pulmonary infection. Our findings indicate that the dynamic regulation of the T6SS is indeed important in the infected lung. While expression of H3-T6SS does not appear to be critical in the initial stages of pulmonary infection, the active selection for loss of function mutants optimized to exploit host itaconate is highly relevant to support persistent infection in patients. Appreciating the complexity of the *P.aeruginosa* adaptation to the lung, it is not surprising that these pathogens thrive in patients despite seemingly potent antibiotics. Antimicrobial susceptibility testing as defined using planktonic growth in artificial media does not accurately predict clinical success against such versatile pathogens that readily assume biofilm and intracellular forms of growth, even under conditions of limited O_2_. Effective eradication of infection, especially pulmonary infection, caused by these pathogens will require novel therapies that consider the metabolic machinery that enables *P. aeruginosa* to maintain viability in diverse environmental conditions.

## Methods

### Key resource tables

**Table 1.** Strains and plasmids used in this study.

| Strain | Relevant characteristics | Source |
| --- | --- | --- |
| *P. aeruginosa* WT PAO1 |  | Riquelme et al. 2020 (1) |
| $\Delta vgrG1$ PAO1 | PAO1 with deletion of PA0091 | This study |
| $\Delta H2$-T6SS PAO1 | PAO1 with deletion of PA1656 — PA1671 | This study |
| $\Delta H3$-T6SS PAO1 | PAO1 with deletion of P2365— PA2372 | This study |
| $\Delta ict$ PAO1 | PAO1 with deletion of PA0883 | Riquelme et al. 2020 (1) |
| $\Delta retS$ | PAO1 with deletion of PA4856 | This study |
| *Escherichia coli* | BW29427 | W. Metcalf, University of Illinois |
| *Saccharomyces cerevisiae* | InvSc1 | Invitrogen |
| pMQ30 | 7.5 kb mobilizable vector *oriT*, *sacB*, GmR | Shanks et al. 2006 (2) |

**Table 2.**
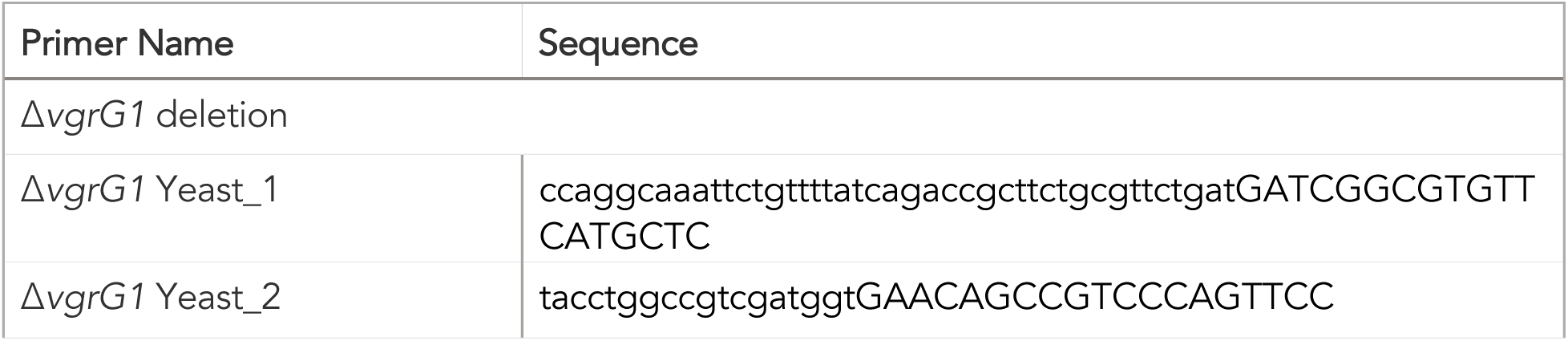
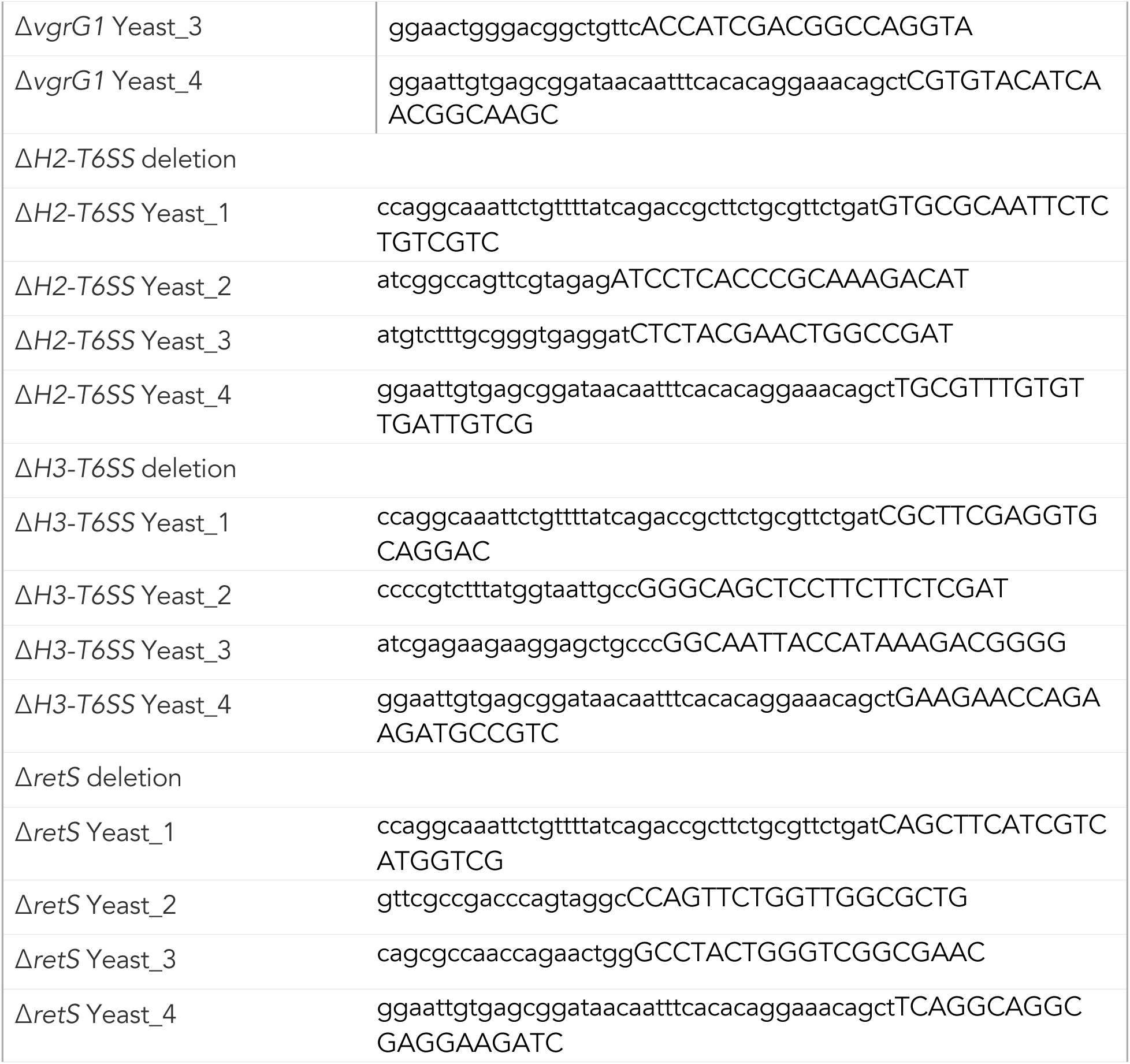
Primers used in cloning.

**Table 3.**
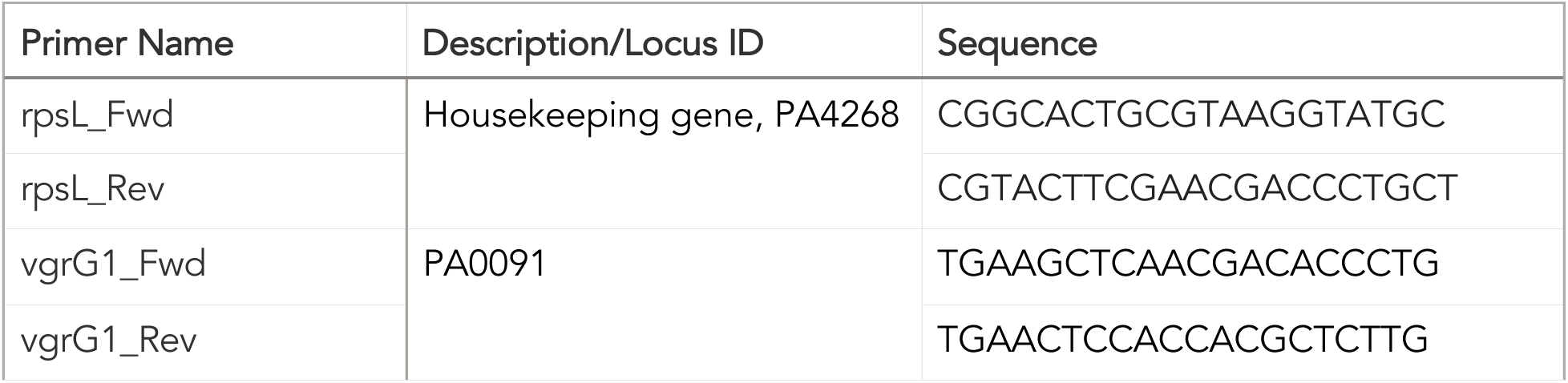
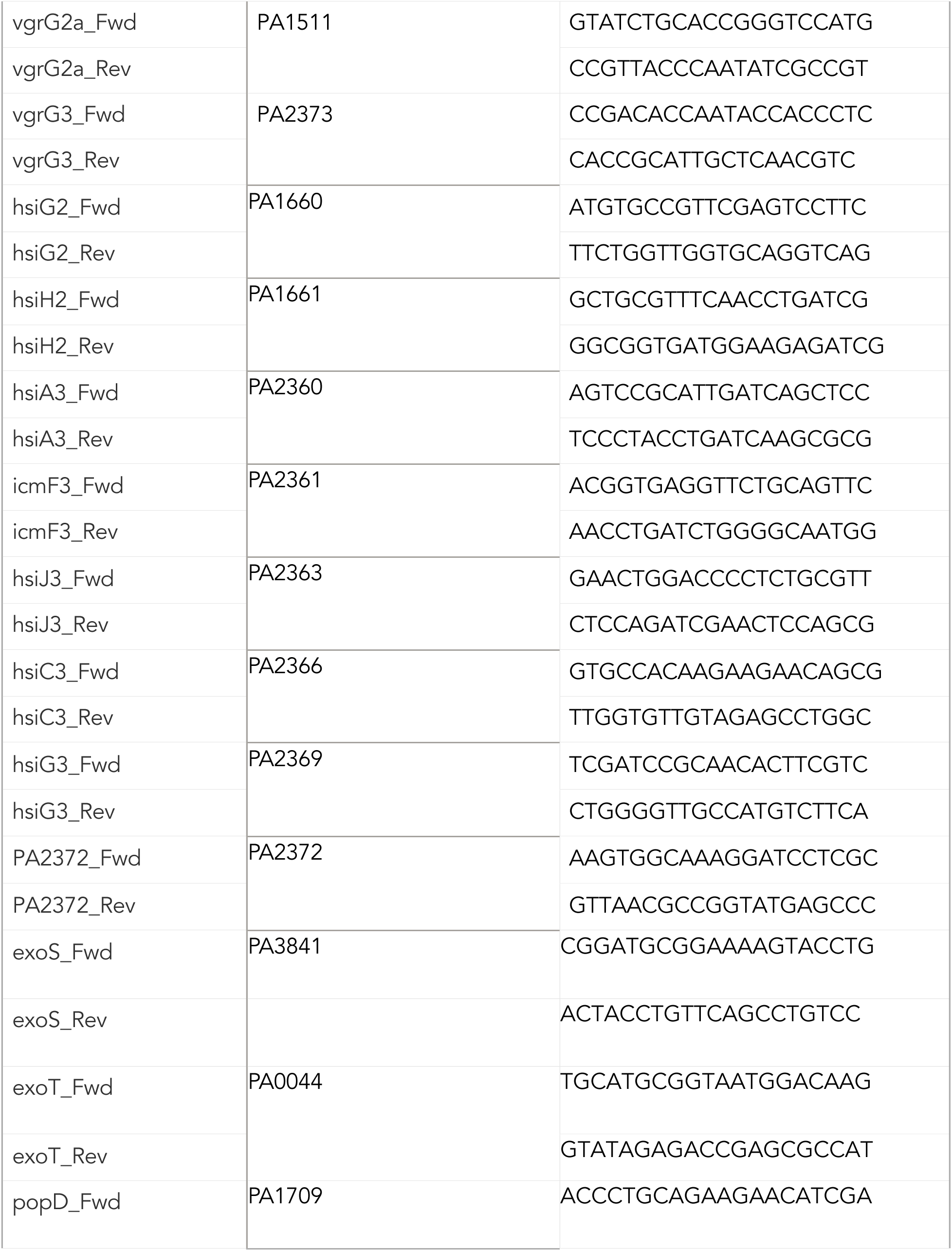
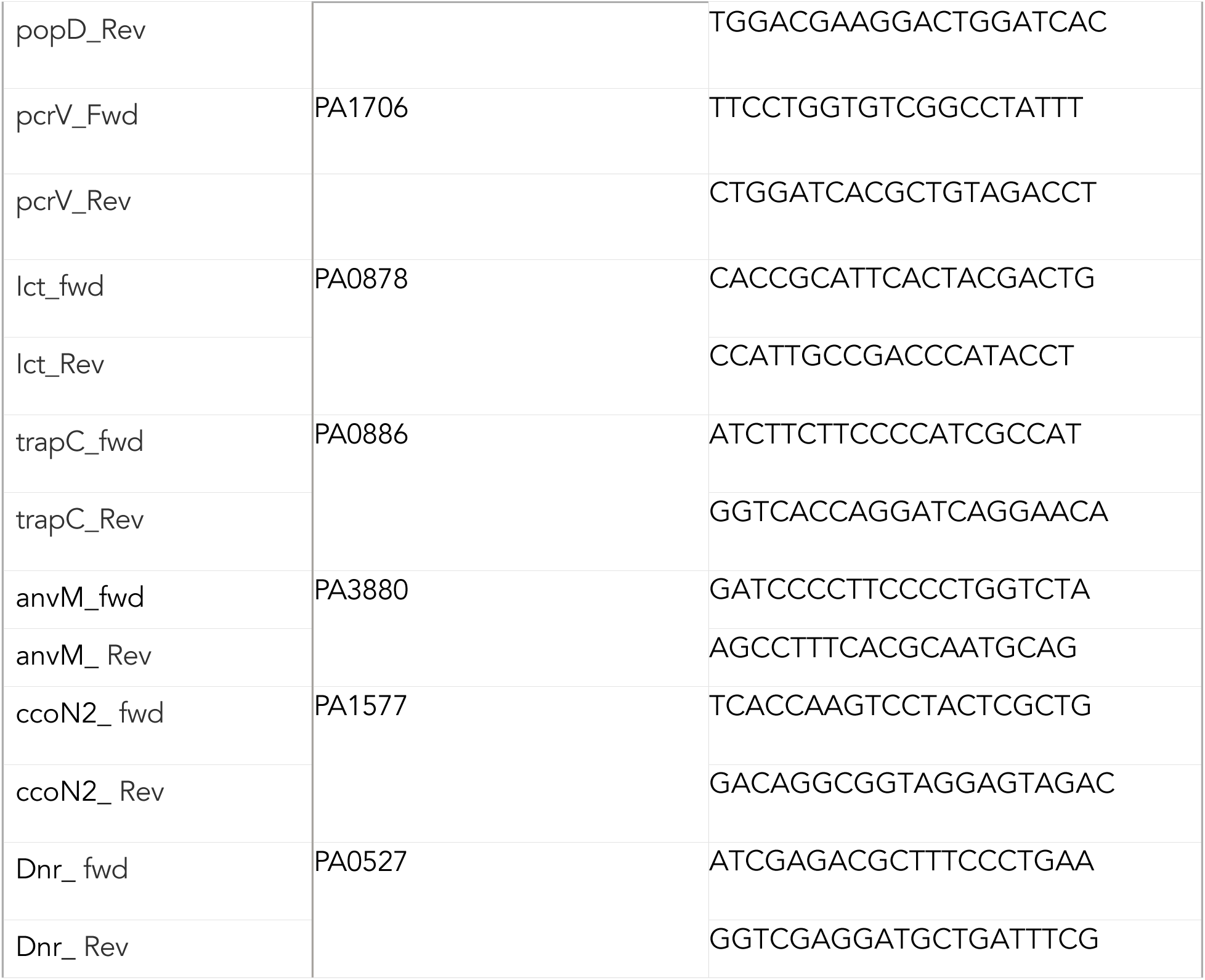
Primers used for qRT-PCR in this study.

### Mouse experiments

All animal procedures were performed in accordance with institutional guidelines at Columbia University Irving Medical Center and were approved under IACUC protocol AABD5602. Wild-type (WT) C57BL/6NJ mice (7–8 weeks old, 20–25 g; Jax #005304) were purchased from the Jackson Laboratory. *Irg1*⁻/⁻ (Acod1⁻/⁻) mice (Jax #029340) were sourced from the Jackson Laboratory and bred on-site at Columbia University Irving Medical Center. Each *in vivo* experiment included equal numbers of male and female mice. Animals were randomly assigned to cages and maintained in barrier facilities under standard housing conditions (12-hour light/dark cycle, temperature 18–23 °C, 30–50% humidity). Mice were provided with standard irradiated chow (Purina Cat #5053, supplied by Fisher).

Flow cytometry for immune cell recruitment in infection studies Immune cell profiling of live cells isolated from bronchoalveolar lavage fluid (BALF) and lung tissue was performed by staining with a fluorescently labeled antibody panel in combination of LIVE/DEAD viability dye (Invitrogen, L23105A). Absolute cell counts were determined using counting beads (10 µL; 15.45 µm Dragon Green beads, Bangs Laboratories Inc., FS07F).The antibody panel included anti-CD45–AF700 (BioLegend, 103127), anti-CD11b–AF594 (BioLegend, 101254), anti- CD11c–BV605 (BioLegend, 117334), anti-SiglecF–AF647 (BD, 562680), anti-MHCII–APC-Cy7 (BioLegend, 107628), anti-Ly6C–BV421 (BioLegend, 128032), and anti-Ly6G–PerCP-Cy5.5 (BioLegend, 127616), each used at a 1:200 dilution in PBS. Cells were stained for 30 min at 4 °C, washed, and subsequently fixed in 2% paraformaldehyde (Electron Microscopy Sciences, 15714-S). Flow cytometric analysis was performed using a BD LSRII flow cytometer with FACSDiva v9 software, and data were analyzed using FlowJo v10. Immune cell populations were defined based on marker expression as follows: alveolar macrophages (CD45⁺ CD11b⁺/⁻ SiglecF⁺ CD11c⁺), neutrophils (CD45⁺ CD11b⁺ SiglecF⁻ MHCII⁻ CD11c⁻ Ly6G⁺ Ly6C⁺/⁻), and monocytes (CD45⁺ CD11b⁺ SiglecF⁻ MHCII⁻ CD11c⁻ Ly6G⁻ Ly6C⁺/⁻).

### Cytokine analysis

Cytokine levels in mouse BALF supernatants were measured by Eve Technologies (Calgary Canada) using bead-based multiplex technology.

### Untargeted metabolomic analysis

High-resolution liquid chromatography–mass spectrometry (LC–MS) metabolite profiling of bronchoalveolar lavage fluid (BALF) samples was performed at the Calgary Metabolomics Research Facility (Calgary, Canada). Metabolites were extracted using a 50% (v/v) methanol (Supelco, #106018) and water solution. LC–MS measurements were conducted on a Q Exactive HF Hybrid Quadrupole-Orbitrap mass spectrometer (Thermo Fisher Scientific) coupled to a Vanquish UHPLC system (Thermo Fisher Scientific). Metabolite separation was achieved using a Syncronis HILIC UHPLC column (2.1 mm × 100 mm, 1.7 µm; Thermo Fisher Scientific) operated with a binary solvent system at a flow rate of 600 µL/min. Mobile phase A consisted of 20 mM ammonium formate (pH 3) prepared in mass-spectrometry–grade water, while mobile phase B consisted of mass-spectrometry–grade acetonitrile containing 0.1% (v/v) formic acid. The gradient elution program was as follows: 0–2 min, 100% B; 2–7 min, 100–80% B; 7–10 min, 80–5% B; 10–12 min, 5% B; 12–13 min, 5–100% B; and 13–15 min, 100% B. A 2-µL aliquot of each sample was injected for analysis. The mass spectrometer was operated in negative ion full-scan mode at a resolution of 240,000, with a scan range of m/z 50–750. Metabolites were identified by matching accurate mass measurements (±10 ppm) and retention times to those of commercial standards (Sigma-Aldrich). Data processing and metabolite annotation were performed using E-Maven version 0.10.0.

### Generation of mutants

*P. aeruginosa* PAO1 strains containing markerless deletions in *vgrG*1, H2-T6SS and H3-T6SS genes were made by homologous recombination(2). Flanking sequence of 1kb was amplified from each side of the target genes (for primers, see Table 2) and were inserted into plasmid pMQ30 using yeast gap repair method in *Saccharomyces cerevisiae* InvSc1. The resulting plasmids were mobilized into PAO1 WT by biparental conjugation using *E. coli* strain BW29427. Markerless mutants arising from double homologous recombination were initially selected on 100 μg ml⁻¹ gentamicin and subsequently counter-selected on 10% NaCl to eliminate the pMQ30 plasmid. Mutants were verified by PCR and sequencing.

### Bacterial RNA extraction from mouse lung

Mice were intranasally inoculated with 1 × 10⁶ CFU of the indicated *P. aeruginosa* strain suspended in 50 μL of PBS. At 16 h post infection, animals were euthanized and lung tissues were harvested for downstream analyses. Total RNA was isolated from lung samples using TRIzol Reagent (Invitrogen) according to the manufacturer’s protocol, followed by DNase treatment with the DNA- free™ DNA Removal Kit (Invitrogen) to eliminate residual genomic DNA. Bacterial RNA was enriched and purified using the MICROBEnrich™ Kit (Invitrogen) in accordance with the manufacturer’s instructions.

### Isolation of bacterial RNA

PAO1 WT, Δ*ict* PAO1, or ΔH3-T6SS strains were grown overnight in LB media. Cultures were diluted 1:100 in LB +/- 20mM itaconate (Sigma, I29204) and incubated at 37 °C until OD₆₀₀ reached 1.0. Approximately 2 × 10⁸ *P. aeruginosa* cells were harvested by centrifugation and resuspended in RNAprotect Bacteria Reagent (QIAGEN, 76506), vortexed at room temperature for 10 min, followed by centrifugation. Pellets were lysed for 10 min at RT in pH 8.0 lysis buffer containing 30 mM Tris (Corning), 1 mM EDTA (Thermo Fisher), 15 mg/mL lysozyme (Sigma), and 200 µg/mL proteinase K (QIAGEN). TRK lysis buffer (Omega Bio-tek, R6834-02) and 70% ethanol were added, and lysates were loaded onto E.Z.N.A. RNA isolation columns (Omega Bio-tek) for RNA extraction according to the manufacturer’s instructions. Residual genomic DNA was removed using the DNA-free™ DNA Removal Kit (Invitrogen, AM1906).

### Complementary DNA (cDNA) synthesis and qRT–PCR

Complementary DNA (cDNA) was synthesized from total RNA using MultiScribe™ Reverse Transcriptase (Applied Biosystems, 43-688-14). Quantitative real-time PCR (qRT-PCR) was performed using gene-specific or housekeeping primers listed Table 3, PowerUp™ SYBR Green Master Mix (Applied Biosystems, 25742), and the StepOnePlus™ Real-Time PCR System (Applied Biosystems). Relative transcript abundance was calculated using the ΔΔCt method.

### Bulk RNA sequencing

Bacterial RNA from PAO1 WT or ΔH3-T6SS strains cultured in LB +/- 20mM itaconate was isolated as described above in Isolation of bacterial RNA section. Ribosomal RNA-depleted cDNA libraries were prepared according to the manufacturer’s protocol using the Universal Prokaryotic RNA-Seq AnyDeplete kit (NuGEN, #0363-32) and sequenced on an Illumina HiSeq platform. Raw base call files were converted to FASTQ format using Bcl2fastq. Quality-filtered reads were aligned to the *P. aeruginosa* PAO1 reference genome (RefSeq: GCF_000006765.1) using STAR aligner v2.7.3a. Aligned reads were processed to assign read group information and identify duplicate reads using Picard Tools v2.22.3. Gene-level raw counts were generated using FeatureCounts from the Subread package v1.6.3. Differential gene expression analysis was performed with DESeq2 in R v3.5.3, and transcripts with fewer than three reads per gene were excluded from downstream analysis. Transcriptional profiles were visualized using volcano plots and heat maps generated in GraphPad Prism (v10.0c).

### Intracellular metabolomics

PAO1 WT or ΔH3-T6SS strains were grown overnight in LB media. Overnight cultures were diluted 1:50 into LB +/- 20 mM itaconate and incubated at 37 °C until reaching an OD₆₀₀ of 1.0. For metabolite extraction, bacterial cultures were pelleted and washed twice with PBS, followed by centrifugation at 2,000 × g for 10 min at 1°C. Bacterial pellets were resuspended in a 3:1 methanol: water extraction solution and lysed through 10 freeze–thaw cycles by alternating immersion in liquid nitrogen and a dry ice–ethanol bath. Cellular debris was removed by centrifugation at 14,000 × g for 5 min at 1°C, and the resulting supernatants were collected and stored for downstream analysis.

Targeted LC–MS analysis was performed using a Q Exactive Orbitrap mass spectrometer (Thermo Scientific) coupled to a Vanquish UPLC system (Thermo Scientific), operated in polarity-switching mode. Metabolite separation was achieved using a Sequant ZIC-HILIC column (2.1 mm × 150 mm, Merck) at a flow rate of 150 μL/min. Mobile phase A consisted of 100% acetonitrile, and mobile phase B consisted of 0.1% NH₄OH and 20 mM ammonium acetate in water. The gradient elution was run from 85% to 30% mobile phase A over 20 min, followed by a wash at 30% A and re-equilibration at 85% A. Metabolites were identified based on accurate mass measurements within 5 ppm and standard retention times. All data were processed and analyzed using MAVEN version 2011.6.17.

### BMDMs infection assay

Bone marrow cells were isolated from mouse femurs and tibias, centrifuged at 500 × g for 6 min, and resuspended in ammonium–chloride–potassium (ACK) lysis buffer for 2 min at room temperature to remove red blood cells. Following lysis buffer neutralization, cells were plated and differentiated in DMEM supplemented with penicillin–streptomycin and 20 ng mL⁻¹ recombinant murine macrophage colony-stimulating factor (rM-CSF; PeproTech) for 5–6 days. Differentiated bone marrow–derived macrophages (BMDMs) were seeded at a density of 0.3 × 10⁶ cells mL⁻¹ in DMEM containing 1% FBS. BMDMs were infected with PAO1 WT, ΔH3-T6SS, PAO1 Δ*ict*, clinical isolate 686 or 686 Δ*ict* at a multiplicity of infection (MOI) of 10 for 60 min, followed by extensive washing to remove extracellular bacteria. After 30 min, gentamicin (50 μg mL⁻¹; Sigma) was added to eliminate remaining extracellular bacteria. Cells were subsequently washed with PBS, detached using TrypLE Express (Life Technologies), serially diluted, and plated on LB agar for bacterial enumeration at 1.5 or 3h post infection. Cell viability was assessed using trypan blue exclusion staining (Life Technologies).

Genomic analysis of clinical isolates:

Publicly available whole-genome sequencing (WGS) datasets of *Pseudomonas aeruginosa* cystic fibrosis (CF) isolates (BioProject accession numbers PRJEB5438, PRJNA528628, PRJNA1023362, and PRJNA1023843) and intensive care unit (ICU) isolates (BioProject accession number PRJNA629475) were retrieved from NCBI and ENA databases for mutation analysis. The reference genome *P. aeruginosa* PAO1 (GenBank accession number AE004091.2) was used as the comparator. Single-nucleotide polymorphisms (SNPs) in *vgrG3* and *clpV3* were identified in CF and ICU isolates using Snippy v4.6.0 (https://github.com/tseemann/snippy), with the PAO1 genome serving as the reference sequence.

### Biofilm quantification

Biofilm formation assays were performed in clear, flat-bottom 96-well plates (Greiner, #M2936) containing M9 minimal media with 0.5% (w/v) glucose +/- 20 mM itaconate. Wells were inoculated with 1 × 10⁸ CFU of *P. aeruginosa* (PAO1 WT or ΔH3-T6SS) and incubated statically at 37 °C for 48 h. Bacterial growth was monitored by measuring OD₆₀₀ using SkanIt Software v7.0 RE. For biofilm quantification, supernatants were removed, wells were washed and air-dried, and biofilms were fixed with 100% methanol and stained with 1% crystal violet. After washing and drying, bound dye was solubilized with 33% acetic acid, and absorbance was measured at 540 nm using a Varioskan Lux plate reader (Thermo Scientific, #3020-82355).

### Growth curves

For bacterial growth curve assays, U-bottom clear 96-well plates (Greiner Bio-One, 650161) were prepared with 198 µL LB or M9 minimal media +/- itaconate as indicated. Wells were inoculated with 2 µL of overnight *P. aeruginosa* cultures (PAO1 WT or ΔH3-T6SS) grown in LB and normalized to OD₆₀₀ = 4. Optical density (OD₆₀₀) was measured every 10 min for 18, 24, or 36 h using a SpectraMax M2 plate reader (Molecular Devices) during incubation at 37 °C with continuous shaking. For microaerobic growth, bacteria were cultured in LB for 24 h in a controlled chamber at 1% O₂. For anaerobic growth, bacteria were cultured statically in LB under 0% O₂ for 22 h at 37 °C, and endpoint OD₆₀₀ was measured. For pH tolerance assays, bacteria were cultured in LB adjusted to pH 5.0 for 36 h.

### THP-1 cell culture and phagocytosis assay

THP-1 cells were stimulated with 1μM Phorbol 12-myristate 13-acetate (Sigma, Cat P15851MG) for 48 hours prior to infection at MOI of 10 for 1 hour. 50 ugmL^-1^gentamicin was added to the media at the desired time point. To analyze intracellular bacterial CFUs, the infected THP-1 cells were gently detached with trypsin (Gibco, Cat 12604021), lysed with 0.05% saponin (Sigma, Cat SAE0073), serially diluted and plated on LB agar at 1.5-and-3-h post infection. THP-1 cell viability was determined by trypan blue exclusion assay(4).

### Confocal microscopy

Infected cells were fixed with 4% paraformaldehyde and permeabilized with 0.05% Saponin. To stain the infected cells, primary antibodies include mouse anti-human LAMP-1 (Proteintech, Cat 65051-1-Ig) or rat anti-mouse LAMP-1 (Proteintech, Cat 65050-1-Ig) and rabbit anti-Pseudomonas (Invitrogen, Cat PA1-73116) were used. Respective secondary antibodies including goat anti-rabbit- AF647 (Invitrogen, Cat A32733), goat anti-rat-AF647 (Invitrogen, Cat A-21247), goat anti-mouse- AF546 (Invitrogen, Cat A-11030) and phalloidin-iFlour 488 dye (Abcam, Cat AB176753) were used. After staining, the coverslips were washed and mounted on slides with ProLong Silver antifade mounting solution with DAPI (Invitrogen, Cat P36962). High resolution images were acquired with Leica Stellaris 8 confocal microscope at high magnification (63x). Regions of interest were generated for each sample and visualized with Fiji/ImageJ software. Intracellular bacteria associated with LAMP-1 marker were counted manually. Average values were calculated across four images, each containing 20 to 30 cells, from three independent experiments.

### Isolation and visualization of infected alveolar macrophages from mouse lungs

BALF samples were collected from C57BL/6NJ mice intranasally inoculated with PBS, PAO1 WT, or ΔH3-T6SS at 16 h post-infection. BALF immune cells were stained as described in the *Flow cytometry for immune cell recruitment in infection studies* section, except that cells were not fixed prior to sorting. Alveolar macrophages (CD45⁺ CD11b^low^ Siglec-F⁺ CD11c⁺ F4/80⁺) were identified and sorted using a 5-laser Aurora cell sorter. Sorted cells were deposited onto glass microscope slides using a cytocentrifuge (cytospin), then fixed with 4% paraformaldehyde and permeabilized with 0.05% saponin. Staining and visualization were performed as described in the *Confocal microscopy* section. Tile scans were acquired using LAS X v4.1.1 software to determine cell counts for each independent experiment.

### Mitochondrial ROS production

BMDMs infected with either WT PAO1 or ΔH3 PAO1 were detached 16hr post infection. The cells are resuspending the cells in media containing 500nM of MitoSOX Red reagent (Invitrogen, Cat M36008) and incubate for 30 minutes. The samples were collected through BD Fortessa Flow Cytometer and data was analyzed by FlowJo^TM^ v10.

### Carbon source assimilation

Carbon sources were prepared in into predetermined wells in the 96 well microplate (Falcon, Cat 353072) and dried overnight in the sterile environment. The overnight bacterial cultures were standardized, washed and inoculated in 100uL of 1x IF-0a inoculation buffer (Biolog, Cat 72268) supplemented +/- 1x Redox Dye Mix A (Biolog, Cat 74221). The plates were incubated at 37°C without light for 24hr to 48hr, and the growth was measured by OD_600_ and the dye absorbance was measured by OD_590_ using the Varioskan Lux microplate reader (ThermoFisher, Cat VLBL0TD0).

### Single-cell RNA sequencing data reprocessing and analysis

Publicly available single-cell RNA sequencing (scRNA-seq) data from a prior study, archived under GEO accession GSE203352, were downloaded from the European Nucleotide Archive (ENA) (5). Briefly, these data were obtained from lung homogenate of WT or *Irg1^-/-^* C57BL/6NJ mice infected with *P. aeruginosa* via the 10X Genomics Chromium platform. Alignment, filtering, barcode counting, and UMI counting were done using the 10X Cell Ranger (v9.0.1) ‘count’ pipeline against the 2024-A GRCm39 reference transcriptome. Ambient mRNA contamination was estimated and corrected using the package SoupX (6) in RStudio (https://posit.co/download/rstudio-desktop/). Further processing and analysis were performed using the Seurat toolkit (https://satijalab.org). Genes expressed in less than three cells were removed, and cell libraries comprising 7.5% or more mitochondrial RNA were expunged. Data matrices were normalized using SCTransform and integrated using the RPCA algorithm. Unbiased clustering was followed by manual annotation using marker genes from publicly available datasets

### Protein structure

The three-dimensional structures of *Pseudomonas aeruginosa* ClpV3 and VgrG3 proteins were predicted using AlphaFold 3. Structural visualization and mapping of protein domains, conserved cysteine residues, and mutations were performed using PyMOL™ version 3.0.4 (Schrödinger, LLC).

### Statistical analysis

Experiments were not conducted in a blinded manner. All statistical analyses and graphing were performed using GraphPad Prism 9. Data are presented as mean ± SEM, assuming normal distribution. For comparisons involving more than two groups, one-way ANOVA with post hoc multiple comparisons was used. For analyses involving two or more groups across time, two-way ANOVA with post hoc testing was applied. Comparisons between two groups were performed using parametric tests (Student’s *t*-test or ANOVA) when data were normally distributed, or nonparametric tests (Mann–Whitney or Kruskal–Wallis) when normality could not be assumed. A two-tailed *P* < 0.05 was considered statistically significant. Specific *P* values, along with the number of independent experiments and replicates, are provided in the figure legends.

## Data Availability

SNP analysis is done on publically available genomes BioProject accession numbers PRJEB5438, PRJNA528628, PRJNA1023362, PRJNA1023843, and PRJNA629475). scRNA seq analysis was done on available dataset with the GEO accession GSE203352, were downloaded from the European Nucleotide Archive (ENA). Bulk RNA seq and intracellular metabolomics data used in the paper are included in the Supplementary information. The whole genome sequencing and SNP results are provided in the source data file. All the source data are provided in this paper.

## Code availability

This work does not include new code.

## Acknowledgements

This work was supported by NIH grants R01HL170129 (A.P.), R35HL135800 (A.P.), and T32DK0076 (G.G.). The Columbia Center for Translational Immunology (CCTI) Flow Cytometry Core is supported by NIH S10OD030282 and provides access to a 5-laser Aurora cell sorter. Imaging was performed at the Columbia Medicine Microscopy Core (MMC) using a Leica Stellaris 8 confocal microscope, supported by NIH S10 grant 1S10OD032447. Metabolomics data were acquired at the Calgary Metabolomics Research Facility (CMRF), University of Calgary, which is supported by the International Microbiome Centre and the Canada Foundation for Innovation (CFI-JELF 34986).

## Authors Contributions

Conceptualization: A.P, T.W.F.L, B.F, A.Z.B. Methodology: A.P, B.F., T.W.F.L, A.Z.B., Y.T.C, S.R. Investigation: B.F, A.Z.B., Y.T.C., T.W.F.L, G.G, A.T, S.S, I.L. Visualization: A.Z.B., Y.T.C. Funding Acquisition: A.P. Project Administration: A.P. Supervision: A.P. Writing—original draft: A.P. Writing— review editing: A.P., A.Z.B.

## Competing interests

The authors declare no competing interests.

## Notes

### Competing Interest Statement

The authors have declared no competing interest.

### Summary of Updates

The author order has been revised, with the corresponding author placed last.

