## Supplemental information for "T6SS mutants exploit itaconate to support infection of phagocytes"

### Figures and legends

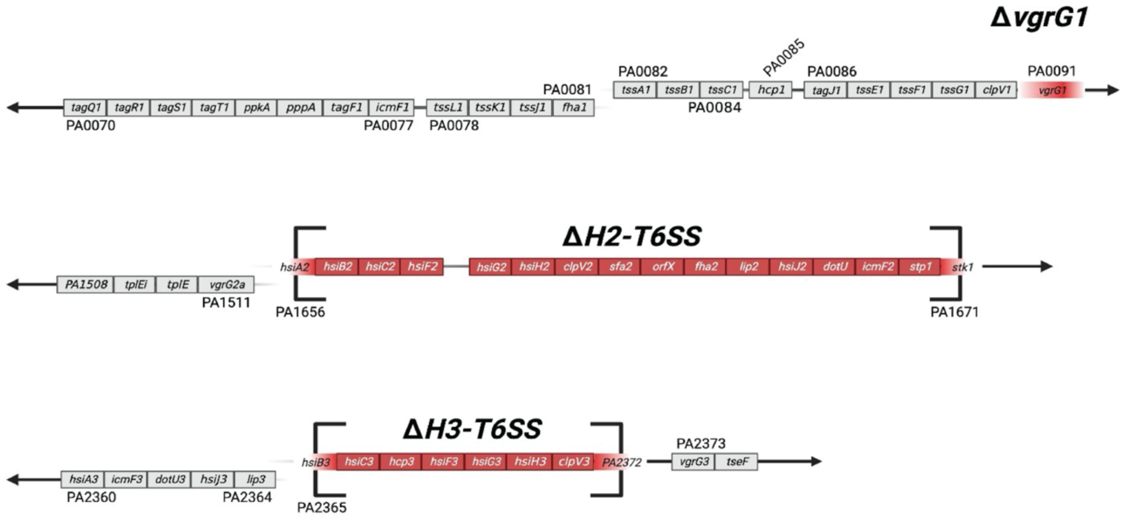


**Supplementary Fig. 1. Deletion of T6SS loci in *P. aeruginosa* PAO1.**
Schematics illustrating deletion of representative gene(s) in each T6SS locus: (**a**) *vgrG1* in H1-T6SS, (**b**) PA1656–PA1671 in H2-T6SS, and (**c**) PA2365–PA2372 in H3-T6SS.

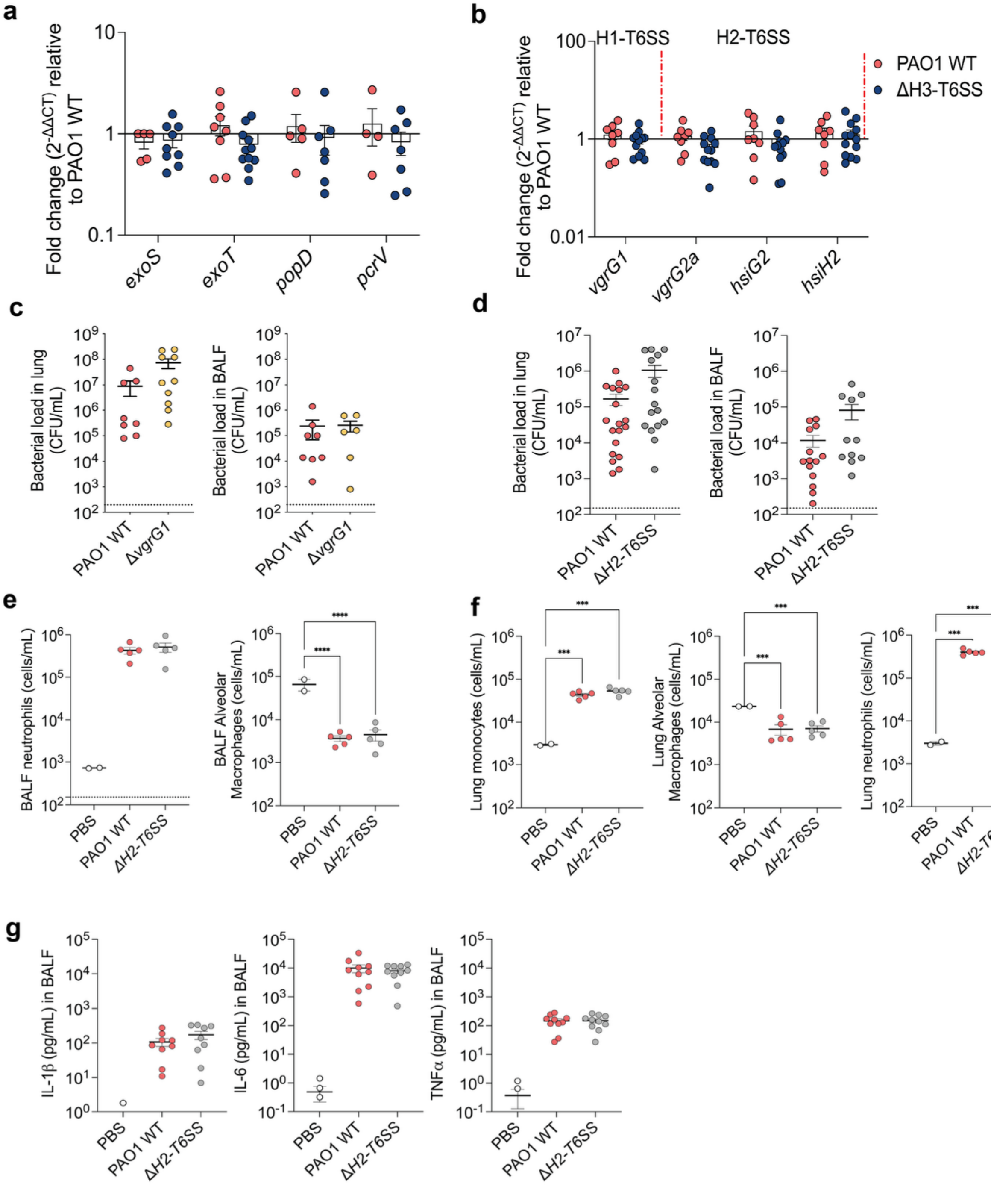


**Supplementary Fig. 2. Effects of the ΔH3-T6SS mutation on in vivo expression of T3/T6 secretion systems (a-b) and roles of H1- and H2-T6SS in pneumonia(c-g).**

(**a**, **b**) In vivo expression of (**a**) T3SS genes and (**b**) H1- and H2-T6SS genes in the lungs of C57BL/6NJ mice infected with ΔH3-T6SS relative to PAO1 WT, as quantified by RT-qPCR. (**c**–**f**) Characteristics of acute pulmonary infection in C57BL/6NJ mice following intranasal inoculation with PAO1 WT, Δ*vgrG1*, or ΔH2-T6SS strains at 16 hpi. Bacterial burden in mice infected with (**c**) Δ*vgrG1* versus PAO1 WT and (**d**) ΔH2-T6SS versus PAO1 WT. (**e–g**) Host immune responses during ΔH2-T6SS or PAO1 WT infection, including immune cell populations in (**e**) BALF and (**f**) lung, and (**g**) cytokine levels in BALF. Data are presented as mean ± s.e.m. Statistical significance was assessed using one-way ANOVA with Tukey's multiple-comparisons test (**e**, **f**). All statistical tests were two-sided. Significance is defined as $P < 0.05$; $*P < 0.01$; $**P < 0.001$; $***P < 0.0001$. Source data are provided as a Supplementary source Data file 2.

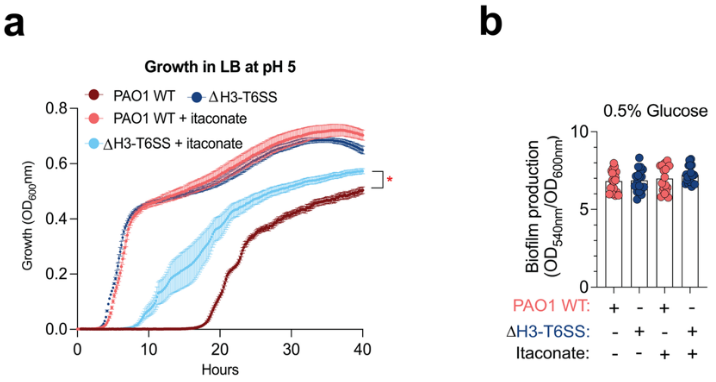


**Supplementary Fig.3. Impact of itaconate on growth characteristics of ΔH3-T6SS and PAO1 WT.**

(**a**) Growth curves of PAO1 WT and ΔH3-T6SS strains cultured in LB +/- itaconate under acidic (pH 5) condition (n = 3). (**b**)Biofilm production by ΔH3-T6SS or PAO1 WT strains grown in M9 media with 0.5% glucose +/- itaconate at 48 h (n = 16). Data are presented as mean ± s.e.m. Statistical significance was assessed by one-way Anova with Tukey's multiple comparisons test (**b**), and Mann Whitney U *t*-test (**a**). Significance is defined as $*P < 0.05$; $**P < 0.01$; $***P < 0.001$; $****P < 0.0001$. All statistical tests are two-sided. Source data are provided as a Supplementary source Data file 3.

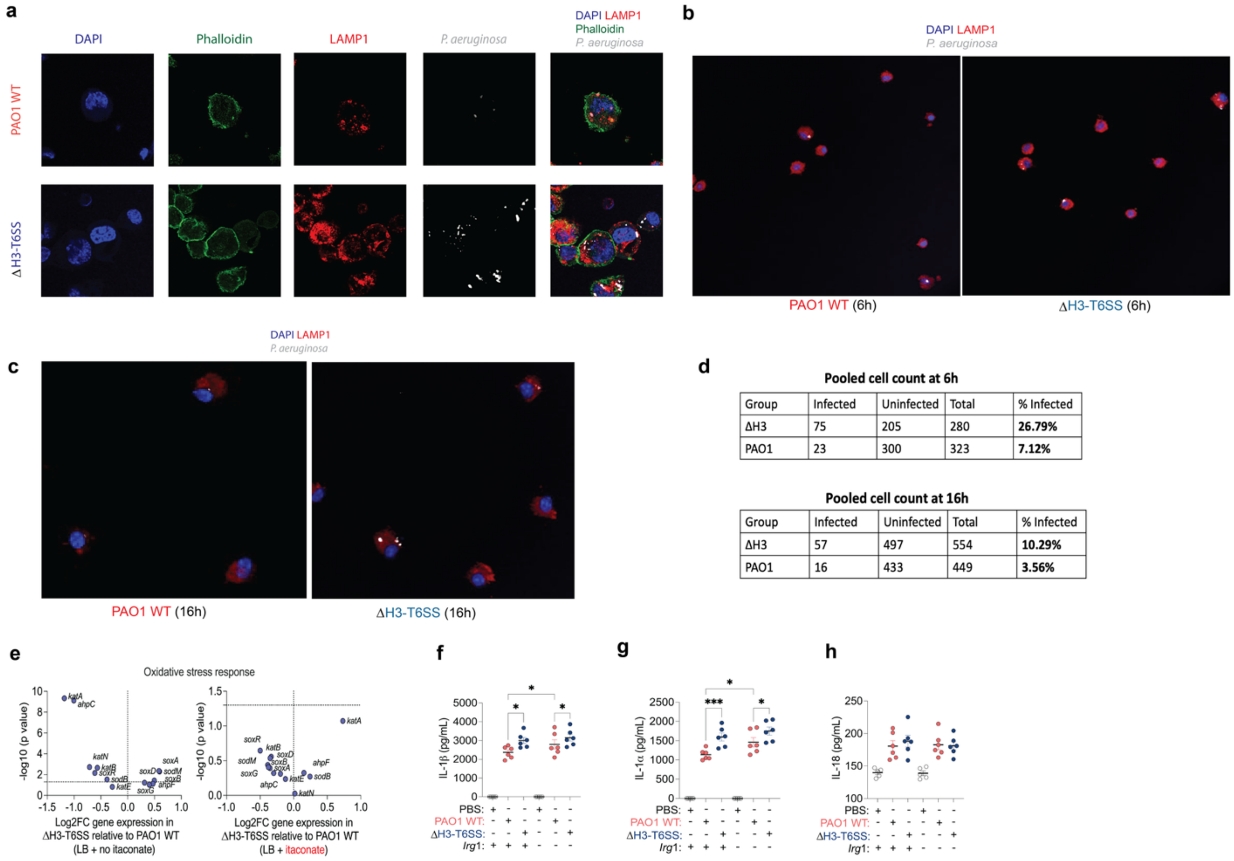


**Supplementary Fig.4. Phagocyte responses to ΔH3-T6SS and PAO1 WT infection.**

(**a**) Confocal microscopy of BMDMs infected with PAO1 WT or ΔH3-T6SS. Cells were stained for nuclei (DAPI, blue), actin (phalloidin, green), lysosomes (LAMP1, red), and bacteria are shown in white. Representative images of infected alveolar macrophages isolated from BALF following infection with PAO1 WT or ΔH3-T6SS at (**b**) 6 hpi and (**c**) 16 hpi. (**d**) Tables of pooled cell counts of infected or uninfected alveolar macrophages at 6 h and 16 h post infection. Expression of genes associated with oxidant stress response in ΔH3-T6SS relative to PAO1 WT grown in LB (**e**) without or (**f**) with itaconate. (**g-i**)**,** Cytokine levels in the supernatants of C57BL/6NJ WT or *Irg1*$^{-}$/$^{-}$ BMDMs infected with PAO1 WT or ΔH3-T6SS: (**g**) IL-1**β**, (**h**) IL-1**α**, and (**i**) IL-18. Data are presented as mean ± s.e.m. Statistical significance was assessed using one-way ANOVA with uncorrected Fisher's LSD test (**g-i**) and Wald *t*-test (**e, f**). Significance is defined as *P < 0.05; **P < 0.01; ***P < 0.001; ****P < 0.0001. All statistical tests were two-sided. Source data are provided as a Supplementary source Data file 4.

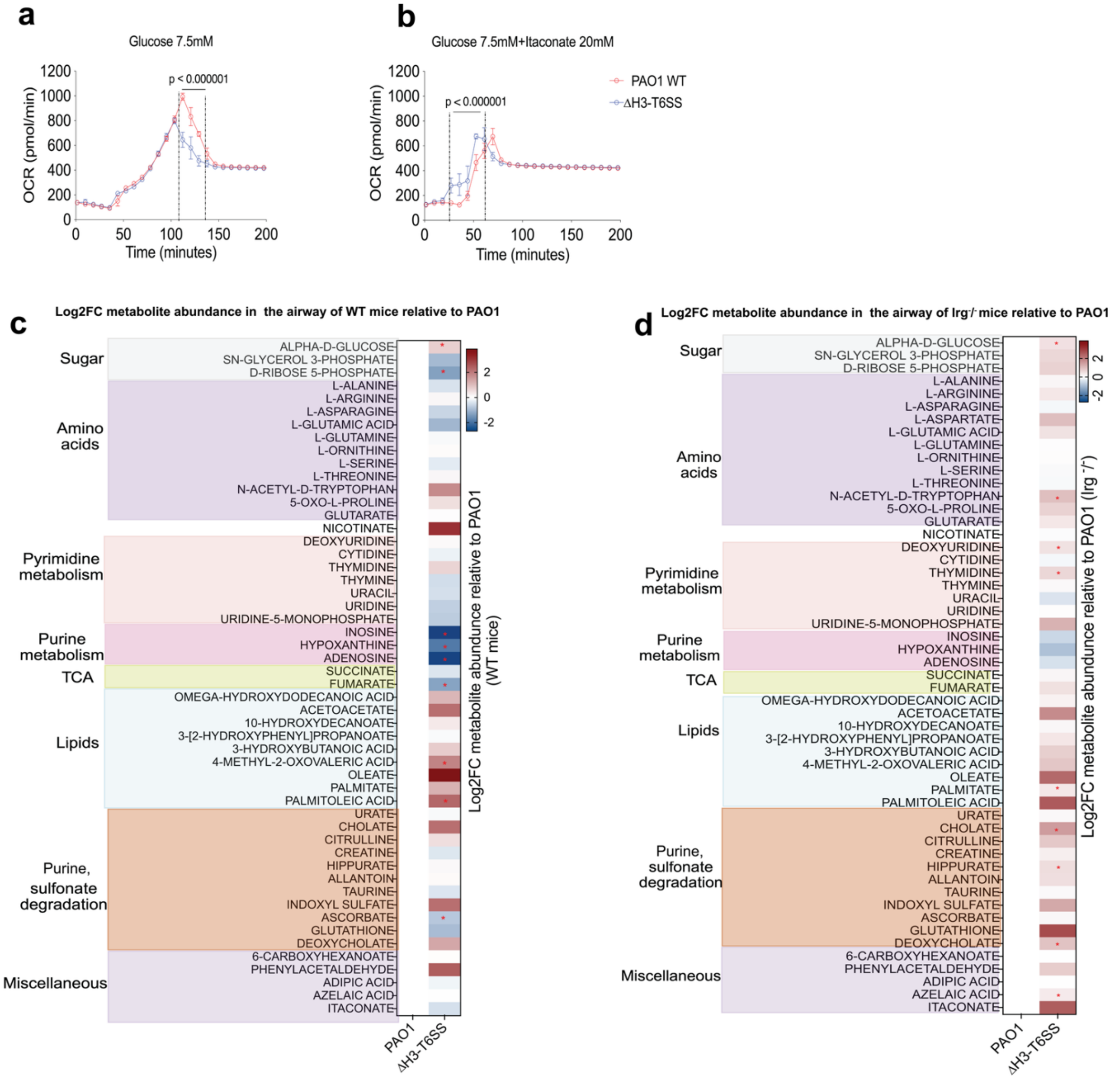


**Supplementary Fig. 5. The ΔH3-T6SS mutation alters *P. aeruginosa* and host airway metabolism.**

Oxygen consumption rate (OCR) of WT vs ΔH3-T6SS PAO1 strains in (**a**) 7.5 mM glucose, (**b**) 7.5 mM glucose + 20 mM itaconate (n = 4-6). Metabolite abundance in the airways of (**c**) C57BL/6NJ WT and (**d**) Irg1$^{-/-}$ mice infected with ΔH3-T6SS or PAO1 WT as shown in the heatmaps (n = 3, with 6 mice in total). Data are presented as mean ± s.e.m. Statistical significance was assessed by unpaired t-test (**a**, **b**) and Student's t-test (**c, d**). Significance is defined as *P < 0.05; **P < 0.01; ***P < 0.001; ****P < 0.0001. All statistical tests are two-sided. Source data are provided as a Supplementary source Data file 5.
